# RNA complexes with nicks and gaps: thermodynamic and kinetic effects of coaxial stacking and dangling ends

**DOI:** 10.1101/2024.04.13.589350

**Authors:** Marco Todisco, Aleksandar Radakovic, Jack W. Szostak

## Abstract

Multiple RNA strands can interact in solution and assume a large variety of configurations dictated by their potential for base pairing. Although duplex formation from two complementary oligonucleotides has been studied in detail, we still lack a systematic characterization of the behavior of higher order complexes. Here we focus on the thermodynamic and kinetic effects of an upstream oligonucleotide on the binding of a downstream oligonucleotide to a common template, as we vary the sequence and structure of the contact interface. We show that coaxial stacking in RNA is well correlated with but much more stabilizing than helix propagation over an analogous intact double helix step (median Δ Δ G_37°C_ ≈ 1.7 kcal/mol). Consequently, approximating coaxial stacking in RNA with the helix propagation term leads to large discrepancies between predictions and our experimentally determined melting temperatures, with an offset of ≈ 10°C. Our kinetic study reveals that the hybridization of the downstream probe oligonucleotide is impaired (lower *k*_*on*_) by the presence of the upstream oligonucleotide, with the thermodynamic stabilization coming entirely from an extended lifetime (lower *k*_*off*_) of the bound downstream oligonucleotide, which can increase from seconds to months. Surprisingly, we show that the effect of nicks is dependent on the length of the stacking oligonucleotides, and we discuss the binding of ultrashort (1 ∼ 4 nt) oligonucleotides that are relevant in the context of the origin of life. The thermodynamic and kinetic data obtained in this work allow for the prediction of the formation and stability of higher order multi-stranded complexes.

**Graphic entry for the Table of Contents (TOC):** 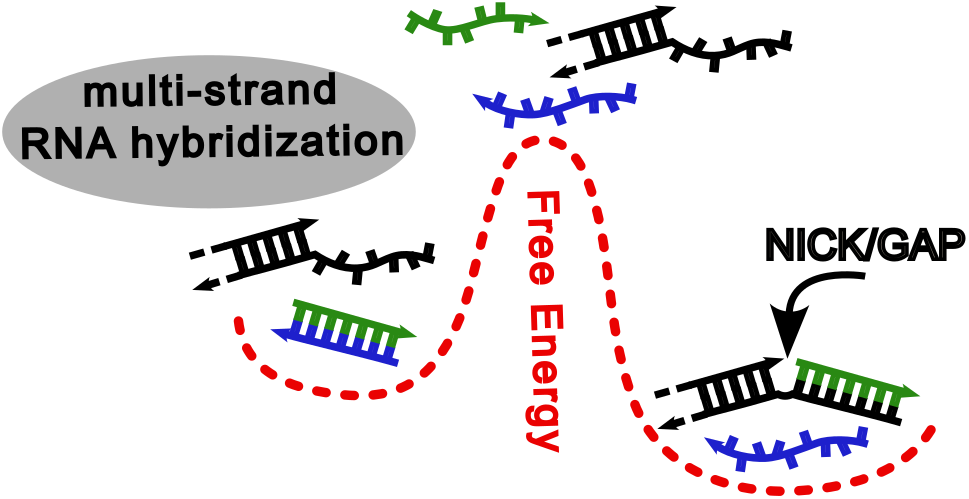

## INTRODUCTION

Understanding the physical interactions among multiple RNA strands is still an open problem. The binding of multiple oligonucleotides to a common template can be cooperative, so that the free energy change due to their hybridization is larger than the sum of their contributions taken individually. The origin of this cooperativity is to be found in the coaxial stacking between the terminal bases of adjacent oligonucleotides. Such stacking interactions have historically been considered to be the driving force in the formation of the double helix^1^, a contributing factor in stabilizing the folding of biologically relevant RNA molecules^2^, and the main driving force for the formation of supramolecular aggregates^3,4^. Systematic studies aimed at quantifying the effect of coaxial stacking in DNA^5,6^ and RNA^7^ through melting experiments date back to the 1990s, highlighting the overall stabilization but a surprising lack of correlation with the energies of helix propagation in DNA. More recently, attempts at assessing the magnitude of base stacking between blunt-ended DNA duplexes in systems without a connecting backbone have been performed both by single-molecule manipulation with DNA origami bundles^8^ and by modeling the behavior of liquid crystalline phases^9–11^, finding comparable values.

To dissect the relative importance of base stacking and hydrogen bonding in the formation of the DNA double helix, the group of Frank-Kamenetskii performed a seminal study in 2006^1^ evaluating the stacked/unstacked equilibrium in nicked duplexes by exploiting their differential electrophoretic mobility. The authors showed that the temperature and salt dependence of coaxial stacking fully explain the temperature and salt dependence of the annealing of two hybridizing strands. Assuming that the helix propagation energies are the sum of base stacking and hydrogen bonding, the authors proposed that base stacking is the main driving force for the propagation of the helix, with hydrogen bonding being either slightly destabilizing (A•T pairs) or negligible (C•G pairs). Opposing this view, experimental studies on single stranded DNA^12,13^ or dinucleotide stacks in solution^14^ and kinetic and thermodynamics of DNA hybridization^13,15^, together with theoretical approaches relying on MD simulations^16^, support the notion that single stranded oligonucleotides can be at least partially pre-stacked in solution, with hydrogen bonding driving the formation of the duplex instead.

Coaxial stacking has been efficiently implemented in predictive tools for RNA hybridization and folding^17–20^, but even the most complete Nearest Neighbor Database (NNDB) of thermodynamic parameters currently available (Turner & Mathews 2004^21^) approximates flush coaxial stacking with the intact helical parameters, making such predictions potentially inaccurate.

Recently, the development of multiplexed single molecule techniques has renewed interest in this topic, leading to updated datasets on coaxial stacking in DNA^22,23^ and yielding free-energy values comparable to those reported from previous melting studies using a short 7 nt long model oligonucleotide^6^. Regarding RNA, the only systematic work performed to date to the best of our knowledge is limited to the characterization of 9 out of 16 possible coaxial interfaces^7^, studied using short 4 nt long model oligonucleotides.

Early work by Pyshnyi & Ivanova showed that coaxial stacking energies in nicked DNA are the same whether the specific experimental model consists of either an oligonucleotide binding to the overhang of a hairpin stem-loop, an oligonucleotide binding downstream of another oligonucleotide on a common template, or an oligonucleotide binding in-between two strands on a common template^24^. This makes the characterization of thermodynamic features in a model system extremely powerful and applicable to a variety of multi-stranded configurations potentially present in high-order nucleic acid complexes.

In this work, we used a combination of fluorescence-based techniques relying on the emission of the adenine analogue 2-aminopurine to provide a systematic study of the thermodynamic and kinetic effects of coaxial stacking and gaps in complexes of multiple RNA strands. Our characterization shows that 7 bp long RNA duplexes are (i) generally stabilized by upstream (towards the 5′ end) dinucleotide gaps, with a behavior analogous to the presence of adjacent unpaired overhangs (3′ dangling ends), and (ii) they are always greatly stabilizing by upstream nicks, with gains in free energy well correlated with helix propagation values over the same sequence in an intact double helix, although much larger.

Furthermore, we dissect the relative contributions of *k*_*on*_ and *k*_*off*_ to the large stabilizations here measured, finding that these entirely result from a slow-down of *k*_*off*_, with *k*_*on*_ being slightly destabilizing and reduced by up to a factor of ∼ 5 by the presence of an upstream oligonucleotide. Finally, we show that the effect of multiple nicks is additive, and is larger with longer stacking oligonucleotides, with implications in the context of the binding of ultrashort oligonucleotides (1 ∼ 4 nt).

Our data is consistent with the idea that single stranded RNA is heavily structured in solution. Building on this notion we disentangle the contributions of base stacking and hydrogen bonding to the formation of the RNA double helix following the approaches of Frank-Kamenetskii^1^ and Zacharias^16^.

## MATERIALS AND METHODS

### General

All measurements in this work were performed in 10mM Tris-HCl and 1M NaCl. Buffer was prepared using a 1M Tris stock solution acquired from Invitrogen and the pH was adjusted using HCl. NaCl powder and concentrated HCl solution were from MilliporeSigma. The concentration of oligonucleotides stock solutions was determined either using a NanoDrop 2000 from Thermo Scientific or a Datrys Ultrospec 2100 pro, and the extinction coefficients computed with the IDT OligoAnalyzer^25^.

### Oligonucleotides synthesis and purification

Reagents and consumables for oligonucleotides synthesis and purification were acquired from ChemGenes and Glen Research. Reagents for cleavage and deprotection were acquired from MilliporeSigma. Oligonucleotides were synthesized on an H-6 K&A solid-phase nucleic acid synthesizer, following the manufacturer recommended protocol. Synthesized, protected oligonucleotides were cleaved from the solid support for 15 minutes at room temperature with a 1:1 v/v mixture of ammonium hydroxide (30% NH_3_ in water) and aqueous methylamine. The nucleobases of the cleaved material were deprotected for 15 minutes at 65 ºC, followed by evaporation of ammonia and methylamine in a vacuum centrifuge and lyophilization of the residual solution. The 2′-OTBDMS protecting groups were removed by dissolving the lyophilized material in 100 μL DMSO and 125 μL TEA.3HF and heating it at 65 ºC for 2.5 hours. After cooling, the deprotected oligonucleotides were precipitated with 0.1 V of ammonium acetate and 5 V of isopropanol for 20 minutes at –80 ºC. The pelleted oligonucleotides were washed once with 80 % ethanol, dissolved in 100 μL neat formamide, and purified by denaturing 20 % PAGE. The desired gel bands produced by the pure oligonucleotides were cut out, crushed, and soaked for 16 hours in a solution of 5 mM sodium acetate pH 5.5 and 2 mM EDTA pH 8. The extracted oligonucleotides were concentrated and desalted using C18 Sep-Pak cartridges (Waters). All sequences used in this work are reported in Supporting Table 1.

**Table 1.**
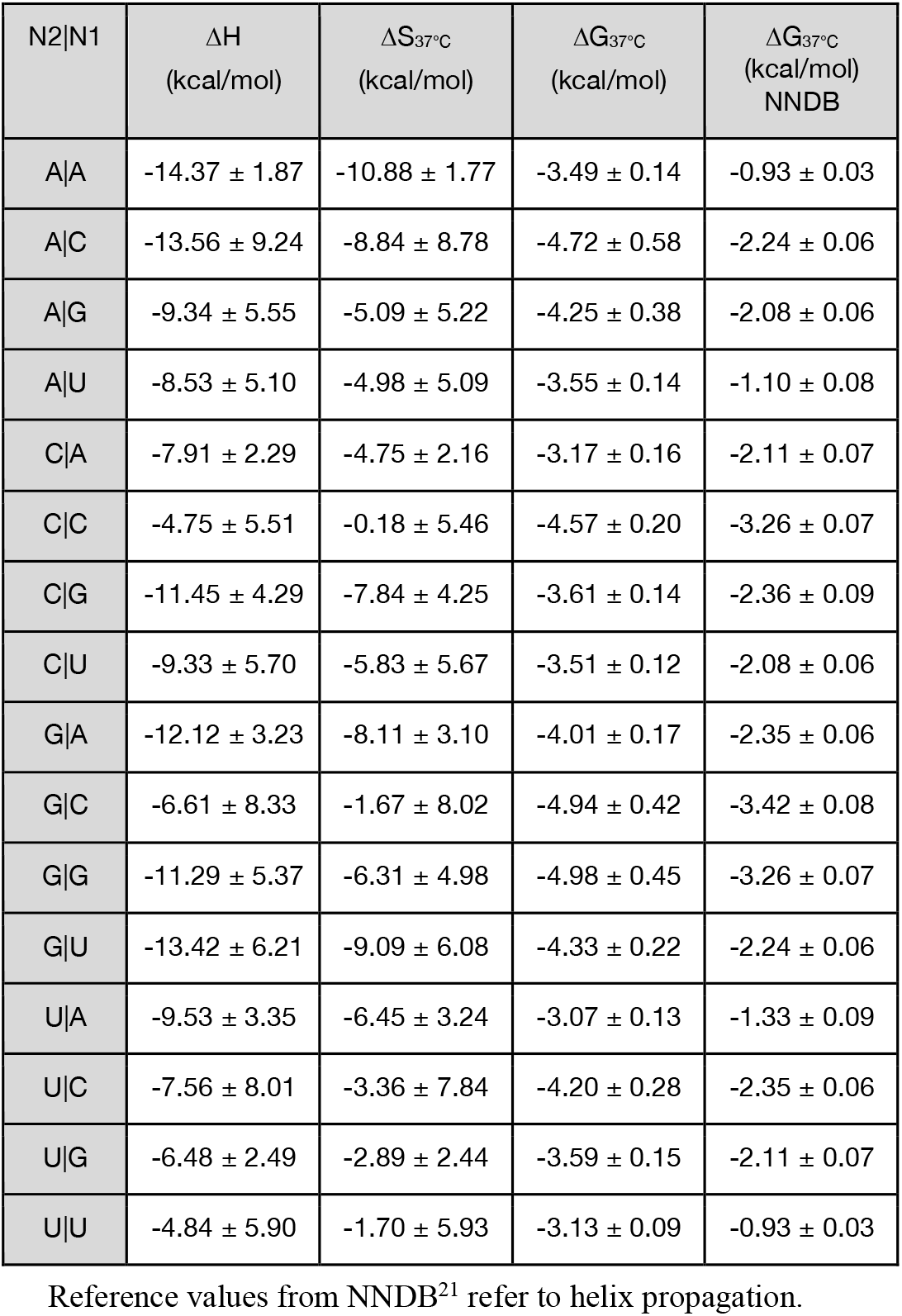
Coaxial stacking energies contributed by upstream nicks in RNA.

| N2\|N1 | $\Delta H$<br>(kcal/mol) | $\Delta S_{37^\circ\text{C}}$<br>(kcal/mol) | $\Delta G_{37^\circ\text{C}}$<br>(kcal/mol) | $\Delta G_{37^\circ\text{C}}$<br>(kcal/mol)<br>NNDB |
| --- | --- | --- | --- | --- |
| A\|A | $-14.37 \pm 1.87$ | $-10.88 \pm 1.77$ | $-3.49 \pm 0.14$ | $-0.93 \pm 0.03$ |
| A\|C | $-13.56 \pm 9.24$ | $-8.84 \pm 8.78$ | $-4.72 \pm 0.58$ | $-2.24 \pm 0.06$ |
| A\|G | $-9.34 \pm 5.55$ | $-5.09 \pm 5.22$ | $-4.25 \pm 0.38$ | $-2.08 \pm 0.06$ |
| A\|U | $-8.53 \pm 5.10$ | $-4.98 \pm 5.09$ | $-3.55 \pm 0.14$ | $-1.10 \pm 0.08$ |
| C\|A | $-7.91 \pm 2.29$ | $-4.75 \pm 2.16$ | $-3.17 \pm 0.16$ | $-2.11 \pm 0.07$ |
| C\|C | $-4.75 \pm 5.51$ | $-0.18 \pm 5.46$ | $-4.57 \pm 0.20$ | $-3.26 \pm 0.07$ |
| C\|G | $-11.45 \pm 4.29$ | $-7.84 \pm 4.25$ | $-3.61 \pm 0.14$ | $-2.36 \pm 0.09$ |
| C\|U | $-9.33 \pm 5.70$ | $-5.83 \pm 5.67$ | $-3.51 \pm 0.12$ | $-2.08 \pm 0.06$ |
| G\|A | $-12.12 \pm 3.23$ | $-8.11 \pm 3.10$ | $-4.01 \pm 0.17$ | $-2.35 \pm 0.06$ |
| G\|C | $-6.61 \pm 8.33$ | $-1.67 \pm 8.02$ | $-4.94 \pm 0.42$ | $-3.42 \pm 0.08$ |
| G\|G | $-11.29 \pm 5.37$ | $-6.31 \pm 4.98$ | $-4.98 \pm 0.45$ | $-3.26 \pm 0.07$ |
| G\|U | $-13.42 \pm 6.21$ | $-9.09 \pm 6.08$ | $-4.33 \pm 0.22$ | $-2.24 \pm 0.06$ |
| U\|A | $-9.53 \pm 3.35$ | $-6.45 \pm 3.24$ | $-3.07 \pm 0.13$ | $-1.33 \pm 0.09$ |
| U\|C | $-7.56 \pm 8.01$ | $-3.36 \pm 7.84$ | $-4.20 \pm 0.28$ | $-2.35 \pm 0.06$ |
| U\|G | $-6.48 \pm 2.49$ | $-2.89 \pm 2.44$ | $-3.59 \pm 0.15$ | $-2.11 \pm 0.07$ |
| U\|U | $-4.84 \pm 5.90$ | $-1.70 \pm 5.93$ | $-3.13 \pm 0.09$ | $-0.93 \pm 0.03$ |

Reference values from NNDB<sup>21</sup> refer to helix propagation.

### Melting experiments

Melting of 2-aminopurine containing probe sequences (O1) mixed with either complementary sequences (O2) or sequences assembling in a nicked complex (O3:O4) or in a gapped complex (O3:O5) was studied using a Jasco FP-8500 Spectrofluorometer equipped with an ETC-815 Peltier temperature-controlled cell holder. The hybridization state of the oligonucleotide as a function of temperature was determined by acquiring the fluorescent emission of 2-aminopurine (2Ap), an adenine base analogue that has negligible impact on RNA thermodynamics^26–28^ and whose fluorescence emission is greatly quenched when in a double-stranded state^29^. The signal of 2Ap was collected at 370nm while exciting at 305 nm using a temperature ramp rate of 1°C/min and continuous stirring.

### Analysis of melting traces

Raw fluorescence traces can be analyzed using a variety of approaches. We found that depending on the specific choice of analysis method, the extrapolated ΔG at room temperature do not typically vary more than ∼ 0.5 kcal/mol (Supporting Figures S1 and S2). We found that individually fitting each experiment and averaging the thermodynamic parameters obtained at different oligonucleotides concentrations provides estimates of ΔH and ΔS for our blunt-ended duplexes that were the closer to NNDB predictions (Supporting Table 2), and thus this is our preferred method for reporting our parameters. Comparisons of the analysis methods are available in the Supporting Information.

In order to individually fit each experiment, we have to link changes in the fluorescent traces to the thermodynamic features underlying the hybridization of O1. For a given case study such as hybridization of O1 and O2 to form the O1:O2 duplex at a given temperature, we can derive the equation for a two-state transition. Given the following definitions:

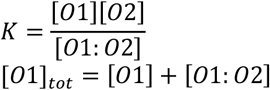

Since the concentration of O1 and O2 are equal by preparation, we can substitute [O2] and [O1:O2] in the first equation to obtain a quadratic equation in terms of [O1]:

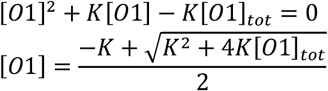

The temperature dependence of hybridization can then be captured within the temperature dependence of *K* as follows:

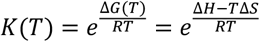

With R being the gas constant (1.987 x 10^−3^ kcal/mol/K) and assuming that ΔH and ΔS are temperature independent in the range of interest, so that the unbound fraction of O1 (*f* = [O1]/[O1]_tot_) in a melting experiment is simply:

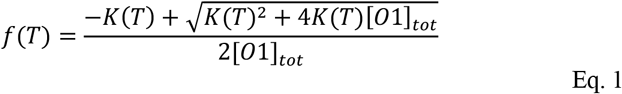

The fluorescent signal coming from our melting oligonucleotides is an average of the signals coming from single stranded O1 oligonucleotides and double stranded O1 oligonucleotides, weighted for their respective emissions, whose temperature dependent behavior can be assumed as being linear for simplicity^30^. To fit our data, instead of manually subtracting baselines from our traces to convert them into fractions, we opted to automatically incorporate them in our fitting routine^31^ for better reproducibility, so that the raw fluorescent signals *(F)* were fitted as:

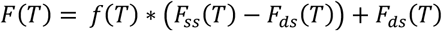

Where *F*_*ss*_*(T) = m*_*ss*_ *+ q*_*ss*_ is the signal coming from single stranded O1 and *F*_*ds*_*(T) = m*_*ds*_ *+ q*_*ds*_ is the signal coming from double stranded O1.

### Error analysis of thermodynamic parameters

Error analysis has been performed following the guidelines established by Turner and colleagues^32^. The coefficients determined from the non-linear fitting of each trace at *N* different total oligonucleotides concentrations using Scipy^33^ have been used to estimate sample mean 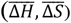, standard deviation (σ_ΔH_, σ_ΔS_) and covariance (σ_ΔHΔS_) for a given sequence under the assumption of normally distributed values with *N*–1 degrees of freedom.

Additionally, the melting temperatures estimated for our oligonucleotide over a 30-fold (or more) concentration range with 6 (or more) samples were fitted to extract alternative estimates of Δ H and Δ S with the following nonlinear equation:

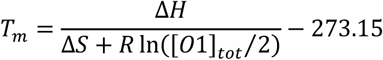

The thermodynamic parameters determined this way have been compared to the individual fits to evaluate the goodness of the assumed two-state transition model (Supporting Figure S1). The coefficients determined from the nonlinear fitting of melting temperatures using Scipy^33^ are provided with their associated standard errors that are typically smaller than the ones deriving from standard deviations of individually fitted traces.

The uncertainty on ΔG at any given temperature was propagated as follows^32^:

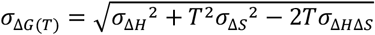

To calculate the stabilization provided by nicks and gaps compared to the control blunt duplexes, uncertainties on each thermodynamic parameter 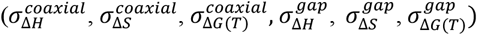 have been independently propagated, so for example for coaxial stacking Δ G:

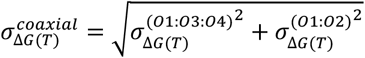

Uncertainties on estimated melting temperatures (in Kelvin units) have been determined as follows:

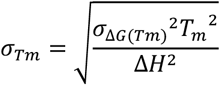

### Kinetic experiments

Hybridization of 2Ap labeled O1 oligonucleotides against complementary strands was studied using a Jasco FP-8500 Spectrofluorometer equipped with an SFS-852T Stopped-Flow accessory. The fluorescent oligonucleo-tide was loaded in a 5mL syringe at a fixed final concentration of ≈ 1μM and the complementary oligonucleotides were loaded in the second 5mL syringe to measure the hybridization process at four different concentrations typically varying from ≈ 3μM to ≈ 0.5μM. Fluorophore was excited at 305 nm and emitted light was collected at 370 nm. Syringes and cell were thermostatted at 20°C for the duration of the experiment.

### Analysis of kinetic traces

The drop of fluorescent signal over time as determined through Stopped Flow experiments was fitted in MATLAB with a bimolecular reaction model numerically integrated using the variable-step solver *ODE15s*, while fixing *k*_*off*_ using our experimentally determined *K = k*_*off*_*/k*_*on*_.

Bimolecular rates and relative errors were determined from the average and standard deviation of the four measurements, weighted for their relative errors as determined by MATLAB *nlinfit*.

## RESULTS AND DISCUSSION

### Experimental Design

To characterize the thermodynamic properties of RNA duplexes in three-strand complexes, we designed a 7 nt long probe sequence (O1) bearing a 5′-phosphate terminus and a 2Ap nucleobase as a reporter for its hybridized/unhybridized state^27^. The 7 nt length was picked to match the melting experiments from Pyshnyi & Ivanova^6^ due to their consistency with more recent single molecule studies^22^.

The probe oligonucleotide was studied either pairing with its perfectly complementary 7 nt oligonucleotide (O2) or to its complementary stretch on a 24 nt long template (O3) bound either to a 17 nt long upstream oligonucleotide (O4) for studies on the effect of nicks or to a shorter 15 nt long upstream oligonucleotide (O5) for studies on the effects of gaps (Figure 1).

**Figure 1.**
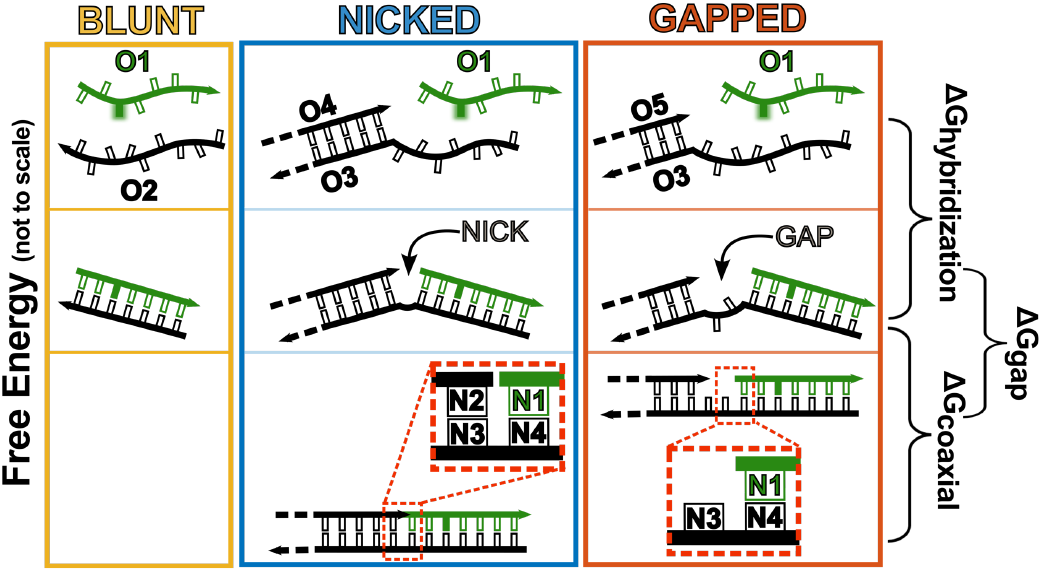
Sequence design. Experimental design to characterize the effect of gaps and nicks on the thermodynamics and kinetics of binding of a short RNA oligonucleotide (O1, green) as part of a blunt-ended duplex (yellow, left box), nicked complex (blue, center box) or a gapped complex (red, right box). 2-aminopurine is represented as a filled green nucleobase in oligonucleotide O1.

Because duplexes separated by single nucleotide gaps are known to interact through coaxial stacking, our design with a dinucleotide gap was chosen to minimize this effect as reported in literature for both DNA and RNA^34–36,1,37^. The 3′ terminal base (N2) of the 17 nt long upstream oligonucleotide, the 5′ terminal base (N1) of the downstream oligonucleotide and the two matching bases of the template (N3 and N4 respectively) were systematically swapped to sample all possible combinations of canonical pairing nucleobases at both nicks and gaps. By comparing the behavior of blunt-ended duplexes (O1:O2) to the same sequences in three-stranded complexes, we could estimate the effect of upstream nicks and gaps on the thermo-dynamics and kinetics of hybridization.

### Thermodynamic effects of nicks and gaps

To dissect the impact of upstream nicks and gaps on binding of the 7 nt long probe, we performed a series of melting experiments analogous to those previously described by Pyshnyi^6^ and Walter^7^. By picking 15 nt long and 17 nt long O4 and O5 oligonucleotides we ensured that as the probe oligonucleotide dissociates, the upstream oligonucleotides remain bound to the template, yielding a two-state transition event. Among all our experiments, we found that only 3 gapped complexes and one nicked complex (N2 = A, N1 = C) deviated from two-state behavior.

For the gapped complexes this is easily explained in terms of interactions between the O3 overhangs of two O3:O5 partial duplexes (Supporting Figure S3), while no immediate explanation to the non-two-state behavior could be found for AC nicks. In almost all cases studied here, the presence of an upstream gap is stabilizing, while in all cases the presence of a nick is greatly stabilizing relative to formation of the O1:O2 blunt duplex.

We started our analysis by comparing the experimentally determined melting temperatures (with [O1] = 1μM) with predictions from the NN model. The predicted melting of the blunt-ended O1:O2 duplexes could be calculated following the guidelines from the NNDB. For the melting of O1 in gapped complexes, since O1 is next to an unpaired stretch (so-called dangling end) we asked whether the gap had any additional effect over the expected 3′ dangling end stabilization. For O1 oligonucleotides in nicked complexes, we attempted to model the stabilization due to coaxial stacking with the helix propagation term for uninterrupted duplexes as recommended in the NNDB, and as implemented in widely used tools for modeling of multi-strand complexes such as NUPACK^17^ and oxRNA^19^ due to the lack of an available coaxial stacking dataset.

As expected, we found that the melting behavior of blunt-ended duplexes is very well predicted, with a mean ΔT_m_ of only 0.4°C between observed and calculated values (Figure 2, yellow symbols). Interestingly, the melting of O1 from gapped complexes is also well captured by assuming that the entire stabilization results from the standard dangling end effect, with mean ΔT_m_ = 1.6°C between observed and calculated values (Figure 2, red symbols). This finding sets an energetic basis for literature observations that a dinucleotide gap is enough to suppress most interactions between two adjacent duplexed regions. In contrast, the assumption that coaxial stacking effect would be comparable to the helix propagation term fails to predict the melting of O1 from nicked complexes, with the experimentally determined melting temperatures being distributed around a mean ΔT_m_ of 9.9°C above the predicted values (Figure 2, blue symbols).

**Figure 2.**
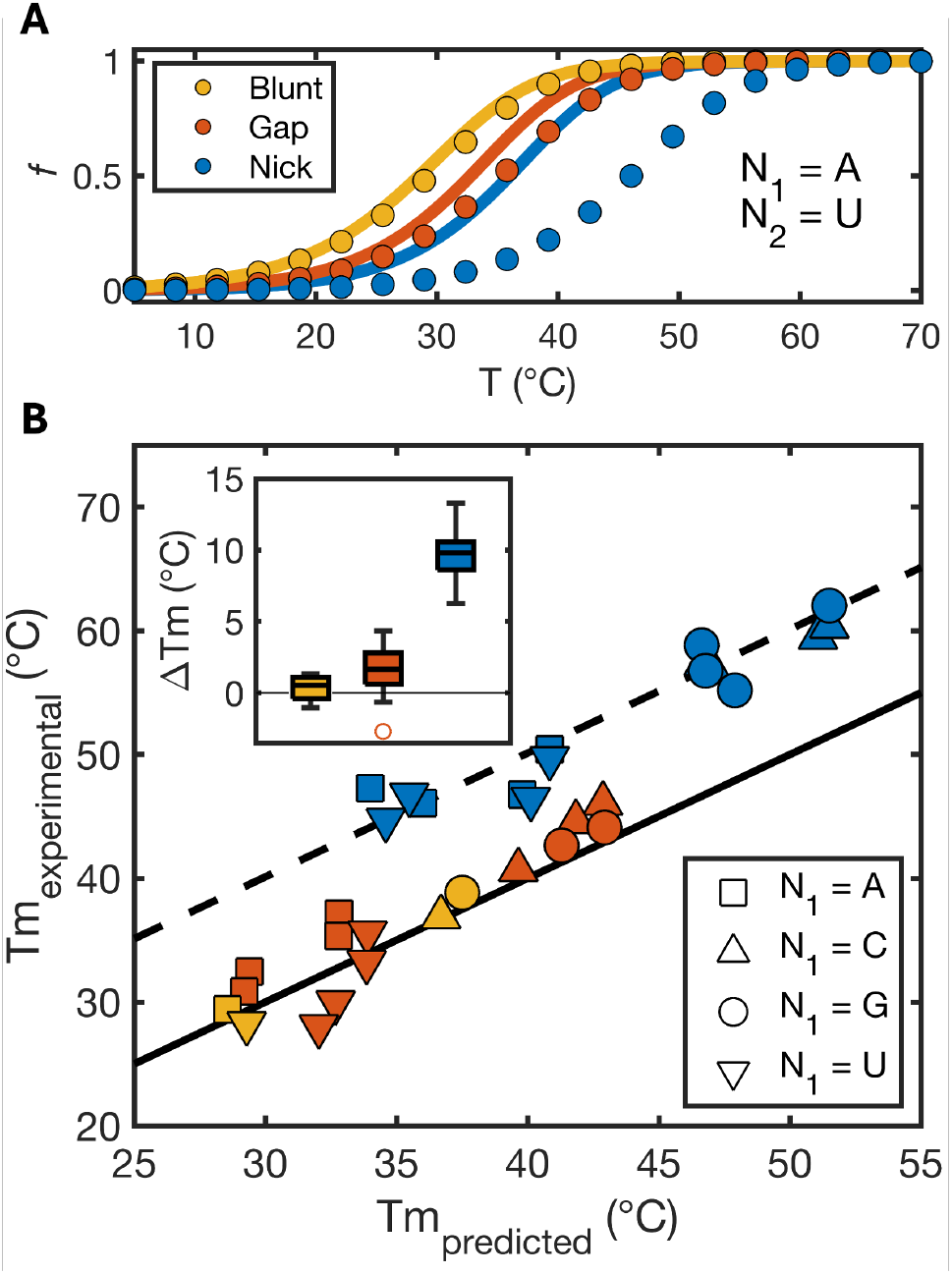
Predicted vs. observed melting of binary and ternary complexes. **(A)** Melting of O1 oligonucleotide (1μM) with N1 = A and N2 = U as part of a nicked complex (blue), gapped complex (red) or blunt-ended duplex (yellow). Scattered dots: fraction unbound calculated from Eq. 1 using experimentally determined values of ΔH and ΔS. Continuous lines were derived using Eq. 1 using predicted values of ΔH and ΔS calculated from NNDB parameters, with the effect of gaps approximated as 3′ dangling ends and the effect of nicks approximated as intact helix propagation energies. **(B)** Overview of melting temperatures of O1 oligonucleotide (1μM) varying N1 and N2. Different symbols refer to different N1 bases in the sequence. Continuous line is the graph bisector (*f(x) = x*) dashed line is the graph bisector shifted by 10°C (*f(x) = x+10*°C). Inset shows a boxplot of differences between measured and predicted melting temperatures. Errors associated with melting temperatures are typically ±0.3°C resulting in error bars that are smaller than markers and thus omitted.

Our thermodynamic study shows that all nicks and most gaps have a stabilizing effect on the binding of the downstream oligonucleotide, relative to formation of a blunt-ended duplex. By looking at differences between the reference blunt-ended duplex and the nicked and gapped duplexes, we could compute thermodynamic effects (ΔH, ΔS, ΔG) of nicks and gaps to directly compare them with literature values for coaxial stacking and 3′ dangling end stabilization (Table 1 for coaxial stacking energies, Supporting Tables 3 and 4 for complete datasets). Given the large uncertainties on ΔH and ΔS determined in this work, which are typical of melting experiments (median σ_ΔH_ = 4.91 kcal/mol, median σ_Δ S,37°C_ = 4.85 kcal/mol), together with their strong dependence on the methodology used to analyze the data (Supporting Tables 2, 3 and 4), comparisons of such parameters with tabulated values may not be reliable. On the other hand, the uncertainties associated with ΔG_37°C_ are very small due to entropy-enthalpy compensatory effects, allowing for more robust comparisons (median σ_Δ G,37°C_ = 0.16 kcal/mol) of values that are less sensitive to the methodology of choice (Supporting Figure S1C). Overall, the comparison of the putative 3′ dangling end stabilization as determined through our experiments with NN database values reveals a remarkable agreement (Figure 3A), with a median absolute difference with reference values of 0.28 kcal/mol and a high degree of correlation, with *r*_ΔG,37°C_ = 0.79.

**Figure 3.**
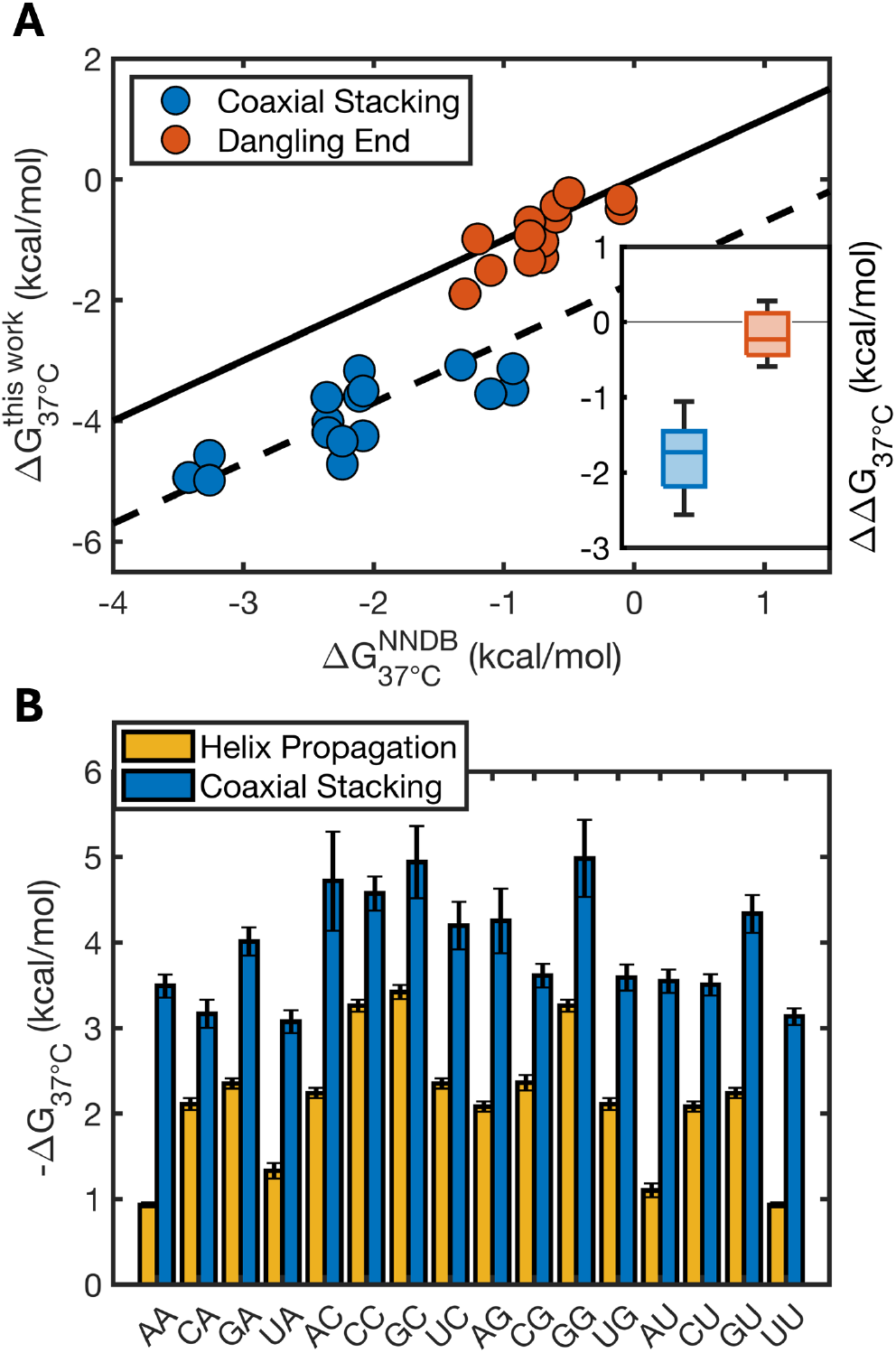
Observed vs predicted stabilization due to nicks (coaxial stacking) and gaps (dangling ends). **(A)** Correlation plot for free energy change due to presence of an upstream gap (dangling end) or upstream nick (coaxial stacking) versus values tabulated in literature. Continuous line is the graph bisector (*f(x) = x*), dashed line is the graph bisector shifted by -1.7 kcal/mol (*f(x) = x-1*.*7 kcal/mol*). Inset shows a boxplot to highlight the distribution of differences between the two sets of values. (**B**) Comparison of free energy change associated with the formation of a new base pair in an intact RNA double helix (yellow) and the energy change for the formation of a coaxial stack at a nicked interface (blue) having the same N2N1 sequence.

Similarly, we found that experimentally determined values ΔG_37°C_ for coaxial stacking and helix propagation values from NNDB are highly correlated (*r*_ΔG,37°C_ = 0.80, Figure 3A,B) as already reported^7^ for RNA but not for DNA^6^. However, these values are significantly offset as expected from our melting study, with coaxial stacking energies always being much greater than their helix propagation counterpart. AA and AC interfaces exhibit the largest differences (Δ ΔG_37°C_ ≈ 2.5 kcal/mol, Figure 3B and Table 1), while the median difference from reference values is -1.7 kcal/mol.

### Kinetic effects of nicks and gaps

Our thermodynamic study revealed that duplexes are significantly stabilized by the presence of an upstream oligonucleotide without or – to a lesser extent – with a gap in the context of three-stranded RNA complexes. Assuming a two-state transition model, this stabilization could reflect either a facilitated hybridization process (higher *k*_*on*_) and/or a prolonged duplex lifetime (lower *k*_*off*_).

To determine the relative contribution of these two possible effects, we performed a series of stopped-flow experiments on our oligonucleotide sets. To our surprise, we found that nicks and gaps always slow down the hybridization of O1 to its complement, with the slowest rate being roughly 5 times lower than the blunt duplex control (Supporting Table 5).

Interestingly, we found that the nicked and gapped configurations exhibit similar decreases in O1 oligonucleotide on-rates, suggesting that a shared mechanism underlies this phenomenon. To test this hypothesis, we measured first the kinetic effect of a dangling 3′ dinucleotide on O1 annealing. Experimentally, this does not significantly affect the hybridization rate of O1, so that even though the magnitude of the stabilization provided by gaps is comparable to the one provided by overhangs, their effects on *k*_*on*_ and *k*_*off*_ are slightly different. Even though the dangling dinucleotide by itself did not result in a significant decrease of the *k*_*on*_, the addition of an extra dangling A_15_ overhang reduced the hybridization rate by roughly a factor of two (Supporting Table 5), suggesting that steric hindrance by the upstream element, whether double or single-stranded, could potentially explain the observed decrease in O1 on-rate in the nicked and gapped complexes and the discrepancy between the effect of an upstream dinucleotide gap and the effect of a simple dangling 3′ dinucleotide.

Given the reduced *k*_*on*_ in the three-strand complexes and large thermodynamic stabilization coming from upstream oligonucleotides, it follows that *k*_*off*_ must be greatly reduced. Indeed, our analysis shows that while blunt-ended O1:O2 duplexes have a characteristic lifetime of 1 to 30 seconds (depending on the identity of N1/N4), the addition of a single coaxial stack can strikingly extend this to up to ∼ 4 months (Figure 4). This result has noteworthy implications for the dynamic behavior of complex mixtures of oligonucleotides, where adjacent oligonucleotides can provide mutual protection from strand dissociation and extend the equilibration timescale for the system even more than previously characterized^27^.

**Figure 4.**
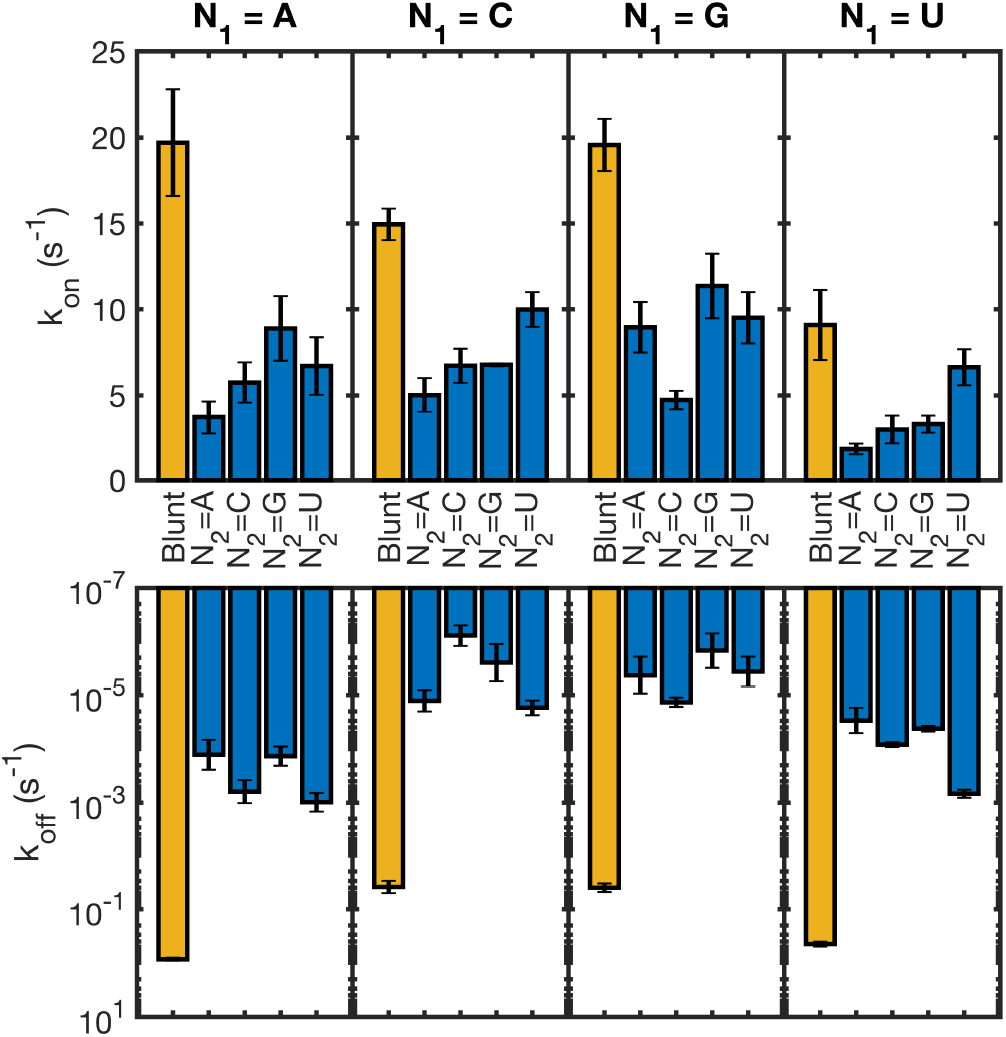
Kinetic effect of coaxial stacking on association (top) and dissociation (bottom) rates. Values of *k*_*on*_ are provided as unimolecular rates at 1μM for easier comparison with *k*_*off*_. Experiments were performed at 20°C.

### Additivity of coaxial stacking

In this work we have characterized the unexpectedly large stabilization arising from coaxial stacking across a single nick site. However, many modern DNA and RNA nanotechnological applications rely on the binding of multiple oligonucleotides in tandem, so that two or more coaxial interfaces stabilize a folded structure^38^; and the nonenzymatic splinted ligation of multiple oligonucleotides has been proposed as a mechanism for ribozyme assembly on the early Earth^39^. The study from Pyshnyi^24^ on DNA tandem complexes revealed that the effect of nicks is cleanly additive, so that the melting of tetramers sandwiched between two adjacent hexamers or octamers could be accurately predicted based on the extrapolated stabilization due to coaxial stacking. We tested the same phenomenon by measuring the melting of one 7 nt long oligonucleotide sandwiched between two 17 nt long oligonucleotides, leading to the presence of an upstream A|U coaxial interface and a downstream U|G coaxial interface.

In our work we have characterized the effect of coaxial stacks for oligonucleotides with an upstream nick. In such system, the unbound state (O3:O4) is not stabilized by a significant dangling end contribution, so that the measured value is close to an effective transition from a reference state with an unstructured terminal to a structured terminal with a novel coaxial stack. When studying tandem oligonucleotides, the process of filling the gap between the two oligonucleotides does not proceed from un unstructured state but from a partially stabilized state due the strong 3′ dangling end contribution of the downstream oligonucleotide (Figure 5A). This makes the effective free energy associated with the transition from the M1:M2:M3 complex to the O1:M1:M2:M3 complex equal to:

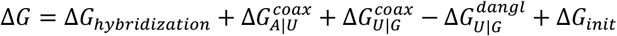

Indeed, this simple model allows us to predict the melting temperature of the oligonucleotide O1 in-between two adjacent oligonucleotides with reasonable accuracy (Figure 5B, compare black circles and solid black line), so that we can conclude that the effects of nicks are additive and that our thermodynamic parameters are appliable to the description of higher-order complexes.

**Figure 5.**
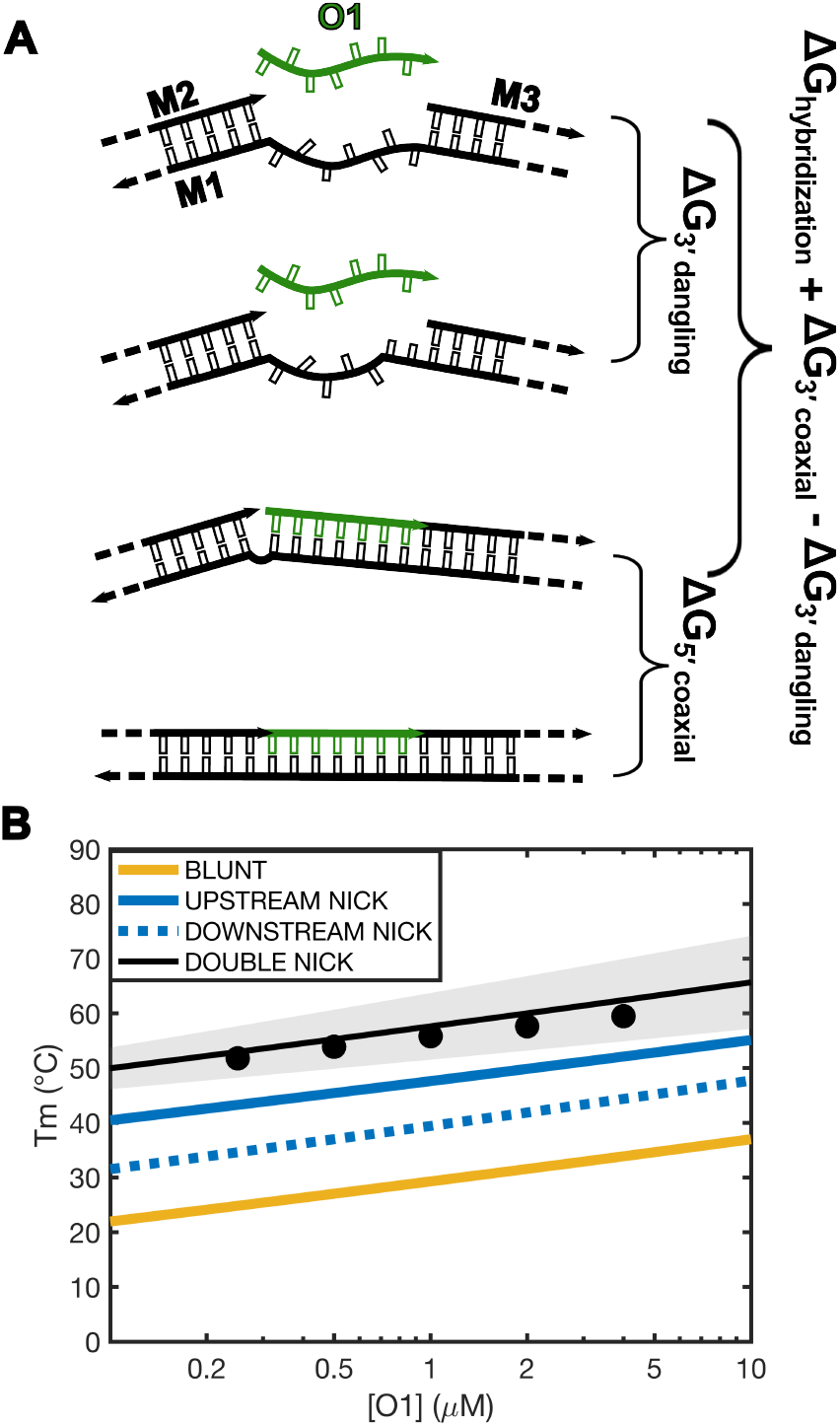
Additivity of coaxial stacking energy for upstream and downstream nicks. The free energy change for O1 filling the gap in-between two adjacent oligonucleotides can be modeled (A) as being the sum of the energy of hybridization of the analogous blunt-ended duplex, together with the contributions of two coaxial stacks and a penalty for the removal of a 3′ dangling end of the downstream oligonucleotide. (B) Melting temperature predicted for O1 in a blunt duplex, with one upstream nick, one downstream nick or both. Scatter points correspond to experimental data for melting of double nicked duplex. Shaded area envelopes the 95% prediction interval for melting temperature of our double nicked oligonucleotide when propagating the errors for coaxial stacking. Lines correspond to predictions for a simple blunt-ended duplex involving O1 (solid yellow) or complexes where O1 has either an upstream (solid blue), downstream (dotted blue) or both (solid black) adjacent oligonucleotide.

### Oligonucleotide length dependence of coaxial stacking energies

Values for helix propagation are typically used as placeholders for coaxial stacking in RNA^21,17,19^. The most comprehensive previous study of the effect of nicks in RNA was performed by Walter & Turner^7^ on 9 out of 16 possible coaxial interfaces in a buffer composition comparable to the one used in this work. Their experimental approach was similar to ours but with a different sequence design that utilized a tetramer oligonucleotide as a probe hybridizing to the overhang of a hairpin stem-loop. Comparing the results from our work with their dataset, we found a remarkably good correlation in coaxial stacking energies (*r*_ΔG,37°C_ = 0.93, Figure 6A). However, we do observe an unexpected median discrepancy of 0.9 kcal/mol between the values of ΔG_37°C_ (Figure 6A, dashed line), with the nicked duplexes measured in our work being systematically more stabilizing. To investigate the origin of this difference, we reduced the length of our 7 nt long O1 to either 5 nt (N2|N1 = A|U) or to 4 nt (N2|N1 = C|C). The nick stabilization measured for the 5 nt long oligonucleotide was comparable to that one measured for its 7 nt long counterpart. This was not true for the 4 nt long oligonucleotide, which we found to be significantly less stabilized by the upstream nick when compared to its 7 nt long counterpart, with Δ ΔG_37°C_ = 0.87 ± 0.16 kcal/mol (Figure 6B). These observations suggest the existence of an interplay between nick stabilization and length of the coaxially stacking oligonucleotides, with a maximum stabilizing effect plateauing at ≈ 5bp. This hypothesis is further corroborated in the next section.

**Figure 6.**
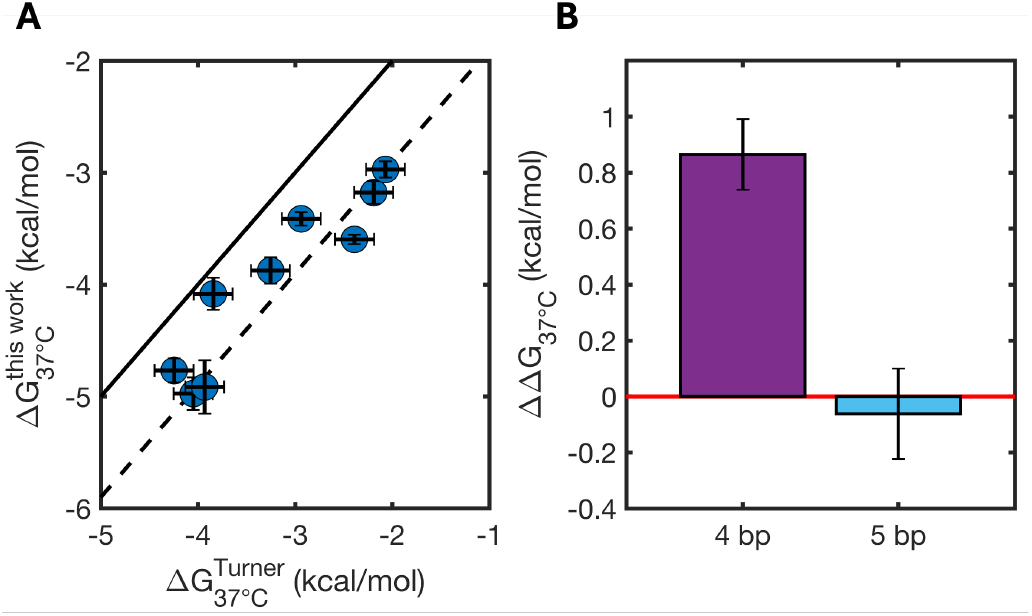
Comparison of coaxial stacking energies with literature data and their length dependence. **(A)** Comparison of stabilization due to upstream nicks as measured in this work and as reported for a limited dataset by Water and Turner. Continuous line represents graph bisector (*f(x) = x*), dashed line is the graph bisector offset by 0.9 kcal/mol (*f(x) = x – 0*.*9 kcal/mol*). **(B)** Difference in coaxial stacking measured in this work with the same value measured when reducing the O1 length from 7 bp down to 4 bp (the same used by Walter and Turner) or 5 bp. Error bars show propagated uncertainties.

### Predicting substrate K_M_ for nonenzymatic primer extension and ligation

Nonenzymatic primer extension and ligation are fundamental chemical reactions believed to have maintained the primordial genetic information within protocells on the early Earth^40^. These reactions rely on binding of chemically activated (5′-phosphorimidazolides) single nucleotides or short oligonucleotides downstream of a primer in a primer-template complex or in the dinucleotide gap between two adjacent oligonucleotides. These substrates have a low intrinsic affinity for their cognate primer/template complexes, but the concentration for half-saturation (K_M_) can be lowered by the presence of a downstream oligonucleotide in a so-called sandwiched system. To date, no reliable tools are available to predict the binding strength of these species since this would require quantitative knowledge of RNA coaxial stacking.

In this section, we aim at formulating predictions for the binding energy of such ultrashort oligonucleotides and to compare them to values estimated from experimental data, assuming that K_M_ is a reasonable estimate of K so that the energies for binding can be approximated as *ln(K*_*M*_*)*RT*. All data discussed here are available in Supporting Table 6.

First, we focused on predicting the binding energy of short oligonucleotides on an overhang (with an upstream nick) as follows,

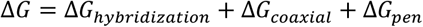

with ΔG of hybridization calculated using the NN helix propagation energies and including an initiation energy penalty of +4.06 kcal/mol at room temperature and extra energetic penalties of +0.63 kcal/mol per terminal AUs^21^. For ΔG_coaxial_ we tested both the predictive power of our newly measured values (Figure 7A) and the predictive power of coaxial stacking values approximated using NNDB helix propagation energies (Figure 7B). For monomers, we replaced ΔG_hybridization_ with the estimated average hydrogen bond energy as previously calculated. We found that the binding strength of short RNA oligonucleotides (down to single monomer) bearing either a 5′-2-aminoimidazole, 5′-methylimidazole or an imidazolium bridged moiety^41,42^ with an upstream nick can be estimated with a high degree of correlation between experimentally derived data and predictions, yielding *r* = 0.97 when using our newly determined coaxial stacking energies, with a mean offset of -0.73 kcal/mol (Figure 7A, circles). Interestingly, the values calculated approximating ΔG_coaxial_ with NNDB helix propagation energies yield an equally high correlation but a much smaller offset, with a mean difference of only + 0.32 kcal/mol (Figure 7B, circles). The discrepancy observed between experimentally determined ΔG and values predicted using our new coaxial stacking energies can be readily explained by the length dependence of coaxial stacking as previously discussed, with a reduced binding energy due to weaker coaxial stacking for short oligonucleotides.

**Figure 7.**
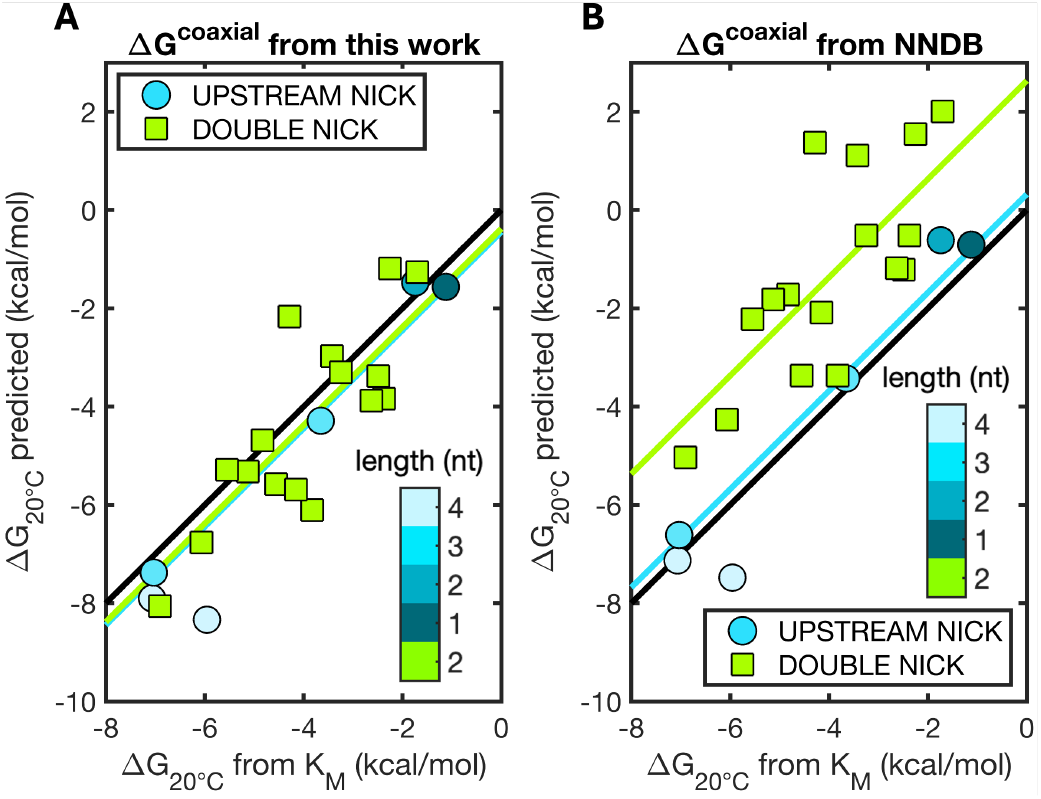
Predictions for binding of short (1 to 4 bp, see colorbar), activated RNA molecules downstream of a primer (circles) or imidazolium bridged dimers in between two adjacent oligonucleotides in a tandem configuration (squares) computed using (**A**) new coaxial stacking values determined in this work or (**B**) coaxial stacking values approximated using helix propagation energies from the NNDB. Black line shows the graph bisector, light blue line shows mean difference between predicted and measured binding energies with upstream nicks, light green line shows mean difference between predicted and measured binding energies with double nicks. Experimental values for comparisons (K_M_) with permission from Zhou et al. (2020)^41^, Ding et al. (2022)^44^ and Ding et al. (2023)^42^. All data are reported in Supporting Table 6.

For double nicked (sandwiched) imidazolium-bridged dinucleotides^43^, we first decided to determine the effect of the chemical activation and the peculiar 5′-5′ linkage to the thermodynamics of hybridization. To do so, we measured the binding of two non-activated dinucleotides (5′-GU-3′ and 5′-UG-3′) to a complementary dinucleotide gap and compared them to their chemically activated counterpart, finding the chemical modification to weaken the interaction on average by 1.55 kcal/mol (Supporting Figure S4).

To calculate the expected binding energy of sandwiched imidazolium-bridged dinucleotides, we assumed that the energies of the two stacking interfaces (nicks) are perfectly additive and penalized by the loss of a dangling end. As previously discussed for tandem systems, the binding energy for a GA dimer in-between two terminal Gs can be computed as follows,

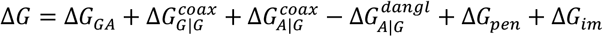

with ΔG_CC_ equal to NN helix propagation energy for a 5′-GA-3′ step and ΔG_im_ equal to the imidazolium-bridge penalty. For ΔG_coaxial_ we tested once again both the predictive power of our newly measured values (Figure 7A) and the predictive power of coaxial stacking values approximated using NNDB helix propagation energies (Figure 7B). For example, using our new values we would have:

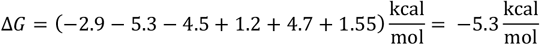

A value extremely close to the experimentally derived one, equal to -5.1 kcal/mol. Relying on our model, we can estimate that the two coaxial stacking terms contribute an astounding 9.9 kcal/mol to the total binding energy, that we would otherwise expect to be equal to +4.5 kcal/mol. This simple example clearly shows how the energy for binding for such short oligonucleotides is expected to come mostly from coaxial stacking, with corresponding binding constants (K) being stabilized by 7 orders of magnitude and effectively dropping from a calculated ∼2000 M (without coaxial stacking) to the experimentally measured ∼100 μM.

When using our newly determined coaxial stacking energies for ΔG_coaxial_ we could obtain values for the binding energies of dinucleotides in double nicked systems (i.e. between two flanking oligonucleotides) in good agreement with experimentally determined values (*r* = 0.82) with a mean offset of -0.38 kcal/mol (Figure 7A, squares) that we understand once again in terms of a reduced binding energy due to weaker coaxial stacking for short oligonucleotides.

By approximating ΔG_coaxial_ with values for helix propagation from the NNDB, we still found a strong correlation (*r* = 0.75) but a larger discrepancy between calculated and experimentally derived energies, with the latter being overestimated on average by +2.63 kcal/mol (Figure 7B, squares).

### Base-pairing and base-stacking contributions to double-stranded RNA formation

Our thermodynamic analysis reveals that the addition of a coaxial stack in an RNA duplex is much more energetically favorable than the formation of a new base-pair in an intact double helix. The hybridization of nucleic acids into a double helix is believed to be driven by the formation of new hydrogen bonds (hb) and stacking interactions. The relative contribution of the two has been a matter of debate, with several experimental results pointing to stacking as being the main contributor^45^. This conclusion has been historically supported by the seminal work by the research group of Frank-Kamenetskii^1^, who performed a quantitative study of nicked DNA duplexes to extract the energy of stacking. Following their approach, the helix propagation term in the formation of a double helix can be described as:

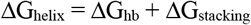

ΔG_stacking_ is assumed to be the energy resulting from coaxial stacking across the nick. One caveat of such analysis is that it treats the formation of a DNA double helix as a transition from a reference state consisting of completely unstructured single strands, so that every new base-pair added to a helix contributes the entire energy of a coaxial stack and two (A•T pairs) or three (G•C pairs) hydrogen bonds, depending on the sequence. Following this interpretation, these authors found that in the transition from unstructured single stranded DNA (with nucleobases bound to water) to the duplexed state (with paired nucleobases), the formation of novel hydrogen bonds between A•T pairs have a net destabilizing effect, and hydrogen bonds between G•C pairs are neutral.

More recently, this interpretation has been challenged by Zacharias^16^, who suggested that treating single stranded oligonucleotides as completely unstructured in solution is not consistent with the strong stacking energies determined from several experimental observations^12–14^. Following the Zacharias approach, the single stranded nucleic acid is treated as partially stacked in solution, so that the free energy change for the duplex formation from these pre-stacked oligonucleotides must be coming mostly from the newly formed hydrogen bonds, with stacking being mildly penalizing.

Applying the approach of Frank-Kamenetskii to our data, the free energy change at 37°C for the formation of hydrogen bonds in RNA would be broadly distributed, yielding an average destabilization of +0.57 ± 0.28 kcal/mol per bond. In contrast, coaxial stacks would be entirely captured by our measurements of nicked duplex stabilization, yielding a ΔG_37°C_ of - 3.96 ± 0.66 kcal/mol. It is however unexpected that the contributions of hydrogen bonds would be so different among different sequences, with coaxial stacks on the contrary being much less sensitive to the sequence context.

Applying the approach by Zacharias on our dataset instead, we found that at 37°C almost all bases (≈ 96%, Supporting Figure S5) are expected to be pre-stacked, and every hydrogen bond in RNA contributes on average -1.04 ± 0.20 kcal/mol, while stacking the remaining (≈ 4%) nucleobases destabilize by +0.06 ± 0.03 kcal/mol (Figure 8) depending on the sequence identity. This analysis provides more readily interpretable results, with hydrogen bond contributions being more narrowly distributed (relative standard deviations equal to 0.20 vs 0.50) and less sequence-dependent, giving further support to the theoretical treatment previously applied to DNA.

**Figure 8.**
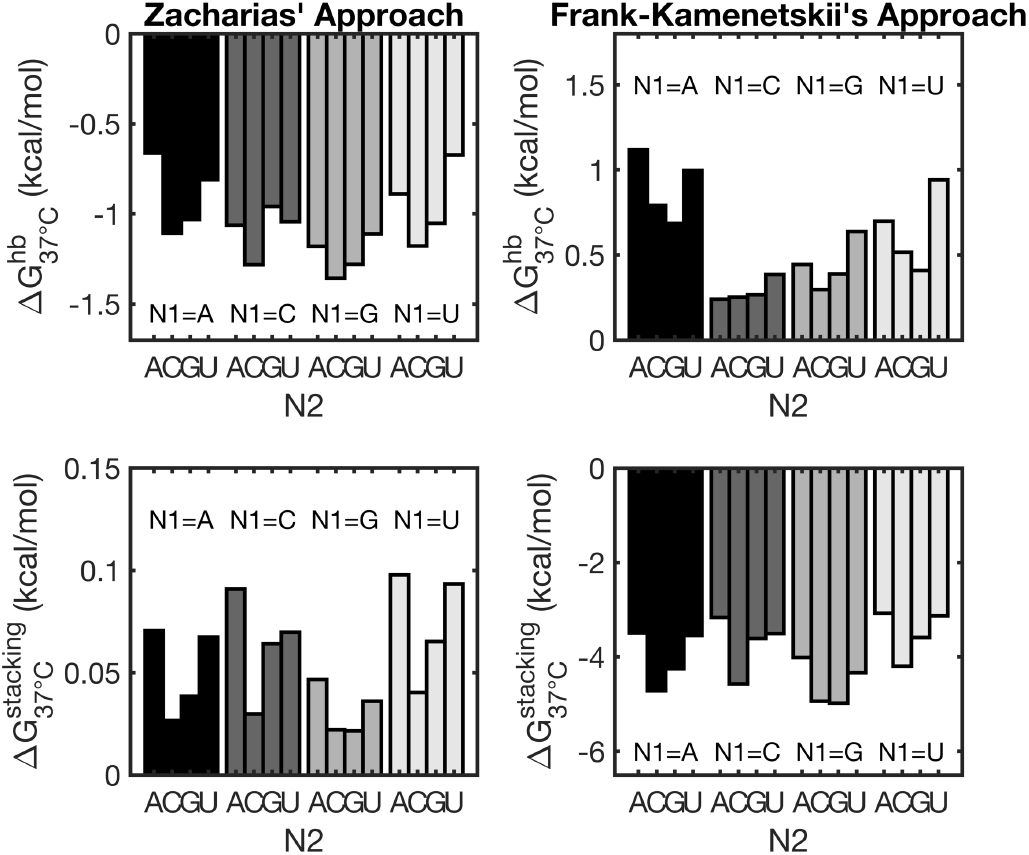
Contributions of hydrogen bonding (top) and stacking (bottom) to the formation of the RNA double helix, using the approaches of Zacharias (left) and Frank-Kamenetskii (right). The approach from Zacharias yields favorable and more tightly distributed energy contributions resulting from hydrogen bonds (- 1.04 ± 0.20 kcal/mol) when compared to the approach from Frank-Kamenetskii (+0.57 ± 0.28 kcal/mol).

## CONCLUSION

In this work we have addressed the challenges arising from the study of three-stranded and four-stranded RNA complexes. The presence of nicks or gaps affects the behavior of a hybridizing strand in a sequence dependent manner. We found that from a thermodynamic point of view, dinucleotide gaps affect oligonucleotide annealing in the same way that simple unpaired stretches (dangling end) would do. This result is in marked contrast with what seen in DNA, where no significant effect coming from upstream dinucleotide gaps has been reported, possibly because energies associated with 3′ dangling ends in DNA are much smaller than in RNA^22^, making gapped oligonucleotides extremely good models for DNA blunt-ended duplexes in surface-tethered setups. In this work we show that this does not hold true for RNA, complicating future thermodynamic and kinetic studies using surface-tethered RNA.

A major conclusion from our work is that approximating coaxial stacking energies with helix propagation energies leads to large discrepancies in the prediction of the melting temperatures of oligonucleotides that bind adjacent to a second oligonucleotide on a common template strand. The coaxial stacking energies as derived from such nicked complexes are well correlated with helix propagation values from the Nearest Neighbor Database, but are much greater than expected. In Table 1 we have presented a complete dataset of energies associated with all 16 coaxially stacking interfaces of canonically pairing nucleobases to improve predictive tools for the thermodynamics of complexes consisting of multiple RNA strands, as well as RNA folding.

A counterintuitive consequence of our findings is that ligating two strands and removing a nick causes a net increase in the total free energy of the system. The origin of this effect has not yet been investigated, but we speculate that the phosphodiester linkage may make optimal stacking geometries inaccessible to the coaxially stacking bases. Alternatively the difference may lie in the loss of conformational entropy for the more constrained ligated structure. We suggest that this may have implications for the energetics of RNA ligase enzymes, and it could pose a thermodynamic explanation for why T4 RNA ligase 2 is more efficient in joining RNA strands over a DNA template (or RNA strands to DNA strands over DNA or RNA templates) with respect to RNA strands on an RNA template in nicked complexes^46^, where a large energy penalty (on top of the energy required to create the new chemical bond) needs to be paid to remove the nick.

We examined the kinetics of oligonucleotide binding and dissociation in order to understand the origin of the thermodynamic stabilization associated with oligonucleotide coaxial stacking. Our data is consistent with steric hindrance by the upstream duplexed region slowing the hybridization (*k*_*on*_) of the downstream oligonucleotide, with the measured thermodynamic stabilization being due to a much greater slowing of the dissociation rate (*k*_*off*_). Importantly, this finding implies that the dynamic behavior of multi-strand complexes is much slower than previously expected^27^, allowing for long lasting metastable states. Such effects are likely to strongly influence models for the nonenzymatic replication of RNA, which is thought to be a critical process for the origin of life. The stabilizing effects that we have observed could, for example, facilitate primer extension by favoring the binding of short substrates such as imidazolium-bridged dinucleotides, adjacent to a primer. On the other hand, the stability of nicked complexes could impede template copying by stabilizing unreactive complexes. Accurate simulation of nonenzymatic RNA replication models must therefore take into account the unexpected stability of nicked multi-strand complexes^47,48^.

Comparing our coaxial stacking energies with the limited literature results, we found our values to be considerably more stable, with a discrepancy deriving from a length dependence of coaxial stacking, setting a physical basis for the unexplained length dependence in the chemical reactivity of imidazolium-bridged oligonucleotides up to 4 bp long^42^. While the origin of this effect is not clear, we speculate that it could possibly be due to the suppression of helix distortions and secondary binding modes – as previously reported for the binding of RNA monomers and dimers in nicked complexes^49,50^ – by longer oligonucleotides.

Moreover, our data provides a quantitative and rational explanation for the strong binding constants measured for extremely short oligonucleotides binding downstream to a primer or in-between two duplexed regions in widely studied nonenzymatic reactions. We found that while all predictions are well-correlated with experimental data, they present significant offsets that can be qualitatively explained in terms of length-dependence and penalty due to the activating imidazolium-bridge moiety, whose effect in the context of a sandwiched (double nicked) system could be estimated as ≈ 1.6 kcal/mol at 20°C.

With this work, we have provided a systematic characterization of the effect of nicks and gaps in RNA, rationalized as deriving from coaxial stacking and dangling end stabilizations. After decades from the first thermodynamic studies applied to nucleic acids, the problem of predicting the rich and complex behavior of multi-strands complexes is still open, and we have devoted our efforts to expand our understanding of such systems. Future efforts should be devoted in implementing our results in predictive tools for thermodynamics of multiple strands, folding and kinetics, and to develop a better understanding of the origin of the length dependence in coaxial stacking and the effect of gaps of different lengths.

## ASSOCIATED CONTENT

### Supporting Information

The Supporting Information is available free of charge on the ACS Publications website.

List of sequences used, comparison of analysis methods, deviation from two-state behavior, full datasets for nicks and gaps thermodynamics and kinetics, study of dinucleotides binding, table with values for short oligonucleotides binding, analysis of hydrogen bonds and stacking contributions in the formation of the double helix (PDF)

## AUTHOR INFORMATION

### Author Contributions

All authors conceived the work. M.T. performed thermodynamic and kinetic study and analyzed results. A.R. synthesized and purified oligonucleotides. All authors wrote the paper, reviewed the results and have given approval to the final version of the manuscript.

### Notes

The authors declare no competing financial interest. Raw data is available at https://doi.org/10.5281/zenodo.10968546.

## ACKNOWLEDGMENT

We thank Mahipal Ganji of IISC Bangalore for initial discussions that prompted us to revisit the topic of stacking energies. We thank members of the Szostak lab for helpful discussions and comments on the paper.

## ABBREVIATIONS

NN: Nearest Neighbor
NNDB: Nearest Neighbor Database
2Ap: 2-aminopurine.

