## Supporting Information for "RNA complexes with nicks and gaps: thermodynamic and kinetic effects of coaxial stacking and dangling ends"

### 1. List of sequences used in this work

The list of oligonucleotides used in this work is provided in Supporting Table 1. Nucleotides N1, N2 (and their complementary nucleotides N4, N3) have been systematically swapped to cover all canonical base-pairs at the interfaces of nicks and gaps.

**Supporting Table 1. Sequences used in this work.**

| Strand type | Sequence | Notes |
|---|---|---|
| O1 | 5′p-**$N_1$**C(2Ap)CGAU-3′ | |
| O2 | 5′-AUCGUG**$N_4$**-3′ | |
| O3 | 5′-AUCGUG**$N_4N_3$**CUGGUACGUGCGACGU-3′ | |
| O4 | 5′-ACGUCGCACGUACCAG**$N_2$**-3′ | |
| O5 | 5′-ACGUCGCACGUACCA-3′ | |
| O2 | 5′-AUCGUG**UG**C-3′ | For kinetic study of dangling ends |
| O2 | 5′-AUCGUG**UG**CAAAAAAAAAAAAAAA-3′ | For kinetic study of polyA tail |
| M1 | 5′-ACUGACCUACGUUGCU**CA**UCGUG**AU**CUGGUACGUGCGACGU-3′ | For additivity study |
| M2 | 5′-ACGUCGCACGUACCAG**A**-3′ | For additivity study |
| M3 | 5′p-**G**AGCAACGUAGGUCAGU-3′ | For additivity study |

Nucleotides in red highlight positions at nicked or gapped interfaces.

### 2. Comparison between analysis methods

In this work, we have characterized the thermodynamics of nicked and gapped oligonucleotide complexes of, with two alternative methods. With one method, each melting experiment was individually fitted, and the coefficients obtained this way were averaged across multiple experiments at different concentrations. Alternatively, the melting temperatures from all experiments were fitted as a function of oligonucleotide concentrations to obtain a second estimate of thermodynamic parameters. Under a two-state description, the values obtained with the two methods should be in agreement.

However, the values for ΔH (Supporting Figure 1A) and ΔS (Supporting Figure 1B) for the two methods are poorly correlated and widely spread. On the contrary, the values for extrapolated ΔG at 37°C (Supporting Figure 1C) are very close (median absolute difference = 0.08 kcal/mol) except for nicked AC complex ($\Delta\Delta G_{37°C}$ = 0.87 kcal/mol).

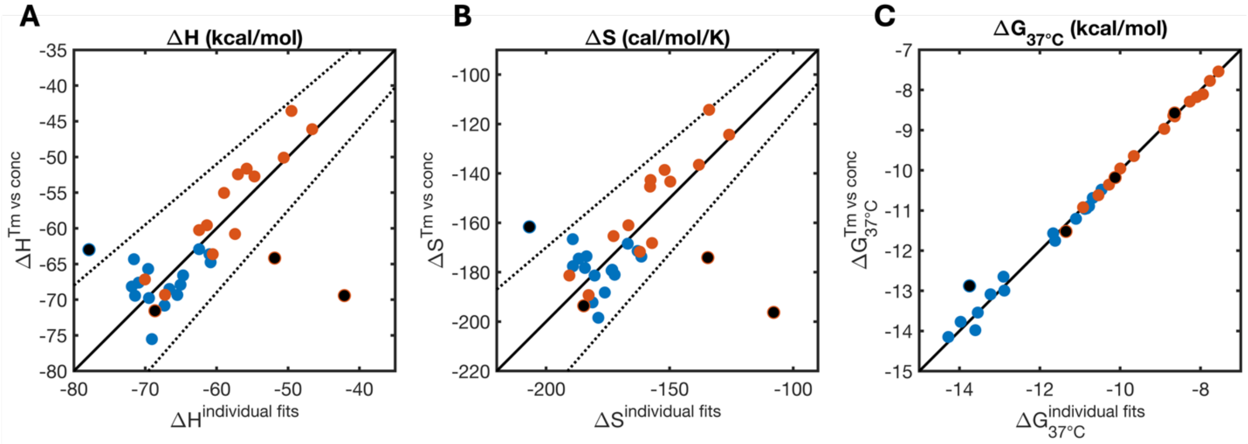

**Supporting Figure S1.** Difference in thermodynamic parameters estimated with two alternative methods as described in the main text. Blue dots referred to nicked complexes, red dots refer to gapped complexes. Black dots refer to anomalous samples that do not follow two-state behavior. Dotted lines show 15% interval bands, considered as a good threshold for establishing adherence to two-state behavior[1,2].

This counterintuitive result derives from compensatory effects in the differences in ΔH and ΔS as immediately visible comparing Supporting Figure 1A and Supporting Figure 1B. The two graphs look extremely similar, so that when the difference is taken to compute ΔG, the discrepancies become extremely small. The further away from the melting temperature at which the thermodynamic parameters have been estimated, the larger the discrepancies between the two methods. We investigated how big of a discrepancy between predictions from the two methods we should expect in a practical range of concentrations and temperatures, as depicted in Supporting Figure 2.

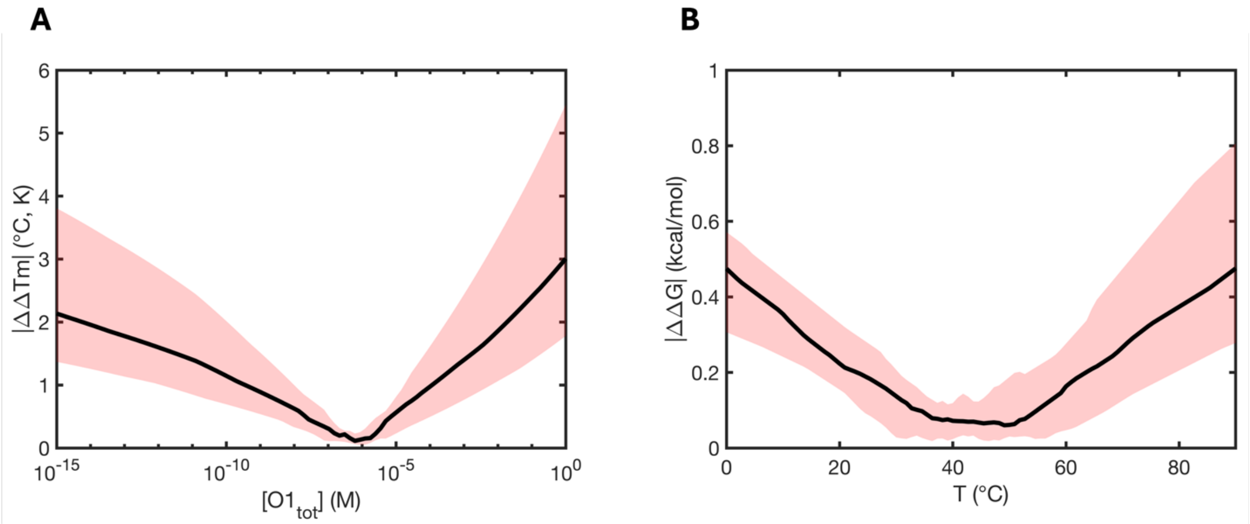

**Supporting Figure S2.** Discrepancies in predictions from the two analysis methods. **(A)** Absolute differences in predicted melting temperatures for our oligonucleotides using the two alternative methods in a practical range of concentrations. **(B)** Absolute differences in extrapolated ΔG as a function of concentration for the two analysis methods. Black lines show median values, shaded areas envelope first and third quartiles for both studies.

We found that discrepancies in extrapolated melting temperatures in a wide range of practical concentrations do not typically vary more than a few degrees Celsius. Similarly, differences in extrapolated ΔG are typically below 1 kcal/mol. The minimum in Supporting Figure 2A aligns with the typical working concentration of this work, while the minimum in Supporting Figure 2B aligns with the typical melting temperatures at our working concentrations, so that in both cases the two methods converge to the same values.

These simple considerations clearly highlight how directly comparing ΔH and ΔS values derived from a limited number of melting experiments can lead to very different conclusions depending on the specific analysis approach picked by the user, and thus in this work we focus on discussing impact of nicks and gaps in terms of ΔG.

Finally, we can compare the values estimated with the two methods for blunt-ended duplexes to the ones computed with tabulated NN values (Supporting Table 2). By doing so, we found that analyzing our traces individually and averaging the fitted coefficients at different O1 concentrations provide values closest to those reported in the NNDB[3], and it is thus our preferred method to report our results for consistency with literature data. The only exception in the context of this work is to be found in the length dependence study, since the lower baseline for tetramers and pentamers could not be reliable estimated given the limitation of our temperature control (lower accessible temperature = -8.0 °C) and we had to rely on Tm vs conc.

### Supporting Table 2. Thermodynamics of blunt ended duplexes

| | NN | | This work, individual fits | | | | This work, Tm vs conc | | | |
|---|---|---|---|---|---|---|---|---|---|---|
| | ΔH | ΔS | ΔH | ΔS | \|ΔΔH\| | \|ΔΔS\| | ΔH | ΔS | \|ΔΔH\| | \|ΔΔS\| |
| **BLUNT, N1=A** | -54.65 | -152.31 | -52.93 | -146.13 | 1.72 | 6.18 | -47.10 | -126.98 | 7.55 | 19.15 |
| **BLUNT, N1=C** | -60.36 | -165.98 | -64.32 | -178.26 | 3.96 | 12.28 | -56.69 | -153.76 | 3.67 | 24.51 |
| **BLUNT, N1=G** | -61.85 | -170.27 | -60.13 | -163.92 | 1.72 | 6.35 | -53.35 | -142.24 | 8.50 | 21.68 |
| **BLUNT, N1=U** | -55.69 | -155.31 | -56.17 | -157.46 | 0.48 | 2.15 | -56.16 | -157.36 | 0.47 | 0.10 |
| **mean** | | | | | **1.97** | **6.74** | | | **5.05** | **16.36** |

Comparison of thermodynamic parameters as determined through the two methods described in this work and compared with reference tabulated NN values for the calculation of blunt ended duplexes thermodynamic features. Values deriving from individual fits are more consistent with predicted features.

### 3. Studying deviation from two-state behavior

The work here presented relies on the assumption that the melting behavior is well described as a two-state transition. We found that this is not true for three case studies involving gapped complexes produced by systematically altering the identity of N1, N2 (and their complementary N4, N3) and for a single case study involving one AC nick. We investigated whether this behavior could be due to self-pairing of two O3:O4 (or O3:O5) duplexes due to their readily available dangling ends. Simple analysis through NUPACK confirms our hypothesis for gapped sequences, but not the nicked sequence (Supporting Figure 3).

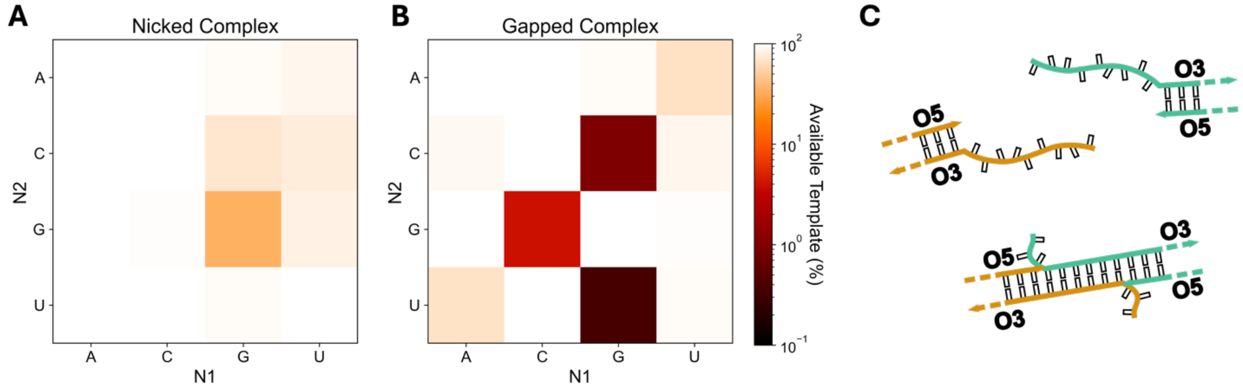

**Supporting Figure S3.** Self-interaction and deviation from two-state behavior. Available template as a function of N1, N2 sequence identity for (**A**) Nicked complexes (O3:O4 duplex), and (**B**) Gapped complexes (O3:O5 duplex). (**C**) CG, GC and UG in Gapped Complexes show signs of strong interaction leading to deviations from the expected two-state transition due to self-interactions. No self-interaction is expected for AC Nicked Complex.

To further test the self-pairing hypothesis, we analyzed the hybridization kinetics: if O3:O4 (or O3:O5) are self-paired, the outcome of the experiment will effectively be due to a relatively slow strand displacement process, and not the much faster conventional nucleation-zipper phenomenon.

Indeed, we find that the three gapped sequences here discussed exhibit an apparent hybridization rate slower by two (GC, $k_{on} \approx 2 \times 10^5$ $M^{-1}s^{-1}$), three (CG, $k_{on} \approx 1 \times 10^4$ $M^{-1}s^{-1}$) or even four orders of magnitude (UG, $k_{on} \approx 6 \times 10^3$ $M^{-1}s^{-1}$) with respect to their relative blunt-ended duplex control (Supporting Table 5).

Once again, AC nicked duplexes do not show signs of self-interaction, with AC gapped duplex behaving in line with other nicked complexes ($k_{on} \approx 5 \times 10^6$ $M^{-1}s^{-1}$, Supporting Table 5), supporting the hypothesis that the deviation from ideal two-state behavior is indeed a feature of this specific nicked interface.

### 4. Thermodynamic parameters for upstream nicks and gaps

In this work we have characterized the thermodynamic effect of an upstream nick or dinucleotide gap on a hybridizing RNA oligonucleotide. Here we report the full dataset of parameters. We have highlighted in red those cases that did not follow a clear two-state behavior and should thus be treated carefully.

**Supporting Table 3. Thermodynamics of upstream nicks (coaxial stacking)**

| N2\|N1 | ΔH individual fits (kcal/mol) | $\Delta S_{37^\circ C}$ individual fits (kcal/mol) | $\Delta G_{37^\circ C}$ individual fits (kcal/mol) | ΔH Tm vs conc (kcal/mol) | $\Delta S_{37^\circ C}$ Tm vs conc (kcal/mol) | $\Delta G_{37^\circ C}$ Tm vs conc (kcal/mol) | $\Delta G_{37^\circ C}$ (kcal/mol) NNDB* |
|---|---|---|---|---|---|---|---|
| A\|A | -14.37 ± 1.87 | -35.07 ± 5.70 | -3.49 ± 0.14 | -23.72 ± 1.87 | -65.24 ± 5.70 | -3.49 ± 0.14 | -0.93 ± 0.03 |
| A\|C | -13.56 ± 9.24 | -28.50 ± 28.32 | -4.72 ± 0.58 | -6.31 ± 9.24 | -7.86 ± 28.32 | -3.87 ± 0.58 | -2.24 ± 0.06 |
| A\|G | -9.34 ± 5.55 | -16.41 ± 16.83 | -4.25 ± 0.38 | -16.43 ± 5.55 | -39.08 ± 16.83 | -4.31 ± 0.38 | -2.08 ± 0.06 |
| A\|U | -8.53 ± 5.10 | -16.06 ± 16.43 | -3.55 ± 0.14 | -10.43 ± 5.10 | -21.99 ± 16.43 | -3.61 ± 0.14 | -1.10 ± 0.08 |
| C\|A | -7.91 ± 2.29 | -15.30 ± 6.98 | -3.17 ± 0.16 | -17.65 ± 2.29 | -46.68 ± 6.98 | -3.18 ± 0.16 | -2.11 ± 0.07 |
| C\|C | -4.75 ± 5.51 | -0.58 ± 17.61 | -4.57 ± 0.20 | -18.82 ± 5.51 | -44.65 ± 17.61 | -4.97 ± 0.20 | -3.26 ± 0.07 |
| C\|G | -11.45 ± 4.29 | -25.28 ± 13.71 | -3.61 ± 0.14 | -10.98 ± 4.29 | -24.41 ± 13.71 | -3.41 ± 0.14 | -2.36 ± 0.09 |
| C\|U | -9.33 ± 5.70 | -18.78 ± 18.28 | -3.51 ± 0.12 | -13.17 ± 5.70 | -30.88 ± 18.28 | -3.60 ± 0.12 | -2.08 ± 0.06 |
| G\|A | -12.12 ± 3.23 | -26.14 ± 10.00 | -4.01 ± 0.17 | -20.81 ± 3.23 | -54.06 ± 10.00 | -4.04 ± 0.17 | -2.35 ± 0.06 |
| G\|C | -6.61 ± 8.33 | -5.38 ± 25.87 | -4.94 ± 0.42 | -10.92 ± 8.33 | -19.83 ± 25.87 | -4.77 ± 0.42 | -3.42 ± 0.08 |
| G\|G | -11.29 ± 5.37 | -20.35 ± 16.07 | -4.98 ± 0.45 | -16.10 ± 5.37 | -36.05 ± 16.07 | -4.91 ± 0.45 | -3.26 ± 0.07 |
| G\|U | -13.42 ± 6.21 | -29.29 ± 19.62 | -4.33 ± 0.22 | -9.53 ± 6.21 | -17.14 ± 19.62 | -4.21 ± 0.22 | -2.24 ± 0.06 |
| U\|A | -9.53 ± 3.35 | -20.81 ± 10.45 | -3.07 ± 0.13 | -15.84 ± 3.35 | -41.48 ± 10.45 | -2.97 ± 0.13 | -1.33 ± 0.09 |
| U\|C | -7.56 ± 8.01 | -10.85 ± 25.28 | -4.20 ± 0.28 | -11.47 ± 8.01 | -23.81 ± 25.28 | -4.08 ± 0.28 | -2.35 ± 0.06 |
| U\|G | -6.48 ± 2.49 | -9.31 ± 7.87 | -3.59 ± 0.15 | -15.15 ± 2.49 | -36.74 ± 7.87 | -3.76 ± 0.15 | -2.11 ± 0.07 |
| U\|U | -4.84 ± 5.90 | -5.49 ± 19.13 | -3.13 ± 0.09 | -7.48 ± 5.90 | -14.03 ± 19.13 | -3.13 ± 0.09 | -0.93 ± 0.03 |

* Values for helix propagation.

**Supporting Table 4. Thermodynamics of upstream gaps (dangling ends)**

| N2\|N1 | ΔH individual fits (kcal/mol) | $\Delta S_{37°C}$ individual fits (kcal/mol) | $\Delta G_{37°C}$ individual fits (kcal/mol) | ΔH Tm vs conc (kcal/mol) | $\Delta S_{37°C}$ Tm vs conc (kcal/mol) | $\Delta G_{37°C}$ Tm vs conc (kcal/mol) | $\Delta G_{37°C}$ (kcal/mol) NNDB* |
|---|---|---|---|---|---|---|---|
| A\|A | -4.08 ± 1.56 | -11.61 ± 4.84 | -0.48 ± 0.12 | -5.33 ± 1.56 | -15.68 ± 4.84 | -0.46 ± 0.12 | -0.10 |
| A\|C | 2.96 ± 5.62 | 11.56 ± 17.98 | -0.63 ± 0.17 | -2.89 ± 5.62 | -7.24 ± 17.98 | -0.64 ± 0.17 | -0.60 |
| A\|G | -0.40 ± 2.50 | 1.88 ± 8.08 | -0.99 ± 0.14 | -10.27 ± 2.50 | -29.51 ± 8.08 | -1.12 ± 0.14 | -1.20 |
| A\|U | 5.57 ± 3.54 | 19.34 ± 11.64 | -0.43 ± 0.08 | 6.06 ± 3.54 | 20.88 ± 11.64 | -0.42 ± 0.08 | -0.60 |
| C\|A | -9.53 ± 5.97 | -26.55 ± 19.70 | -1.29 ± 0.19 | -13.15 ± 5.97 | -38.36 ± 19.70 | -1.25 ± 0.19 | -0.70 |
| C\|C | -5.72 ± 7.44 | -12.34 ± 23.77 | -1.89 ± 0.19 | -10.47 ± 7.44 | -27.59 ± 23.77 | -1.91 ± 0.19 | -1.30 |
| C\|G | -8.51 ± 11.55 | -20.82 ± 36.55 | -2.06 ± 0.28 | -18.20 ± 11.55 | -51.33 ± 36.55 | -2.29 ± 0.28 | -1.70 |
| C\|U | -1.24 ± 3.47 | 0.33 ± 11.42 | -1.34 ± 0.09 | -4.63 ± 3.47 | -10.81 ± 11.42 | -1.28 ± 0.09 | -0.80 |
| G\|A | 3.42 ± 2.92 | 12.09 ± 9.26 | -0.33 ± 0.10 | 3.56 ± 2.92 | 12.73 ± 9.26 | -0.39 ± 0.10 | -0.10 |
| G\|C | 22.21 ± 5.68 | 70.35 ± 18.13 | 0.39 ± 0.33 | -12.75 ± 5.68 | -42.49 ± 18.13 | 0.43 ± 0.33 | -0.40 |
| G\|G | 1.16 ± 2.21 | 6.01 ± 7.32 | -0.70 ± 0.16 | -1.68 ± 2.21 | -3.12 ± 7.32 | -0.72 ± 0.16 | -0.80 |
| G\|U | 9.56 ± 3.94 | 31.54 ± 12.90 | -0.22 ± 0.08 | 10.05 ± 3.94 | 33.00 ± 12.90 | -0.19 ± 0.08 | -0.50 |
| U\|A | -2.85 ± 2.32 | -5.88 ± 7.28 | -1.03 ± 0.11 | -4.55 ± 2.32 | -11.62 ± 7.28 | -0.94 ± 0.11 | -0.70 |
| U\|C | -2.91 ± 8.24 | -4.51 ± 26.21 | -1.51 ± 0.20 | -12.62 ± 8.24 | -35.50 ± 26.21 | -1.61 ± 0.20 | -1.10 |
| U\|G | 8.27 ± 11.97 | 29.35 ± 38.00 | -0.83 ± 0.38 | -10.81 ± 11.97 | -31.82 ± 38.00 | -0.94 ± 0.38 | -1.70 |
| U\|U | 1.47 ± 3.63 | 7.73 ± 11.88 | -0.93 ± 0.08 | 3.43 ± 3.63 | 14.07 ± 11.88 | -0.93 ± 0.08 | -0.80 |

* Values for 3′ dangling end stabilization. No associated uncertainties are available.

### 5. Kinetic effect of upstream nicks and gaps

In this work we have characterized the kinetic effect of an upstream nick or dinucleotide gap on a hybridizing RNA oligonucleotide. Here we report the full dataset of parameters. We have highlighted in red those cases that did not follow a clear two-state behavior. For gapped sequences, this is understood as due to self-pairing of the O3:O5 complexes as discussed in Supporting Data 2. The kinetic rates for such cases have to be interpreted as strand displacement rates and not hybridization rates.
It is important to notice how the deviation from two-state behavior does not create any anomaly in the hybridization rate of C|A, supporting the hypothesis that such deviation is indeed an intrinsic property of that nicked interface.
We have highlighted in light blue the two control experiments (compare with Blunt, N1 = A) for the effect of 3′ a dinucleotide dangling end and an extra polyA$_{15}$ tail.

**Supporting Table 5. Kinetics of upstream nicks and gaps**

| **Experiment (N2\|N1)** | $k_{on}$ ($M^{-1}s^{-1}$) | $k_{off}$ ($s^{-1}$) |
|---|---|---|
| **Blunt, N1 = A** | (1.97 ± 0.31) x $10^{7}$ | $(8.54^{+0.59}_{-0.55})$ x $10^{-1}$ |
| **Blunt, N1 = C** | (1.49 ± 0.09) x $10^{7}$ | $(3.87^{+1.18}_{-0.91})$ x $10^{-2}$ |
| **Blunt, N1 = G** | (1.96 ± 0.15) x $10^{7}$ | $(4.00^{+0.82}_{-0.68})$ x $10^{-2}$ |
| **Blunt, N1 = U** | (0.91 ± 0.20) x $10^{7}$ | $(4.47^{+0.50}_{-0.45})$ x $10^{-1}$ |
| **gapped A\|A** | (0.49 ± 0.08) x $10^{7}$ | $(6.09^{+0.41}_{-0.39})$ x $10^{-2}$ |
| **gapped A\|C** | (0.73 ± 0.08) x $10^{7}$ | $(5.40^{+0.77}_{-0.68})$ x $10^{-3}$ |
| **gapped A\|G** | (0.63 ± 0.13) x $10^{7}$ | $(7.93^{+1.49}_{-1.25})$ x $10^{-4}$ |
| **gapped A\|U** | (0.36 ± 0.08) x $10^{7}$ | $(1.23^{+0.07}_{-0.07})$ x $10^{-1}$ |
| **gapped C\|A** | (0.35 ± 0.09) x $10^{7}$ | $(5.85^{+0.95}_{-0.82})$ x $10^{-3}$ |
| **gapped C\|C** | (0.38 ± 0.07) x $10^{7}$ | $(1.37^{+0.61}_{-0.42})$ x $10^{-4}$ |
| **gapped C\|G** | (1.13 ± 0.11) x $10^{4}$ | $(1.02^{+0.32}_{-0.24})$ x $10^{-7}$ |
| **gapped C\|U** | (3.03 ± 0.38) x $10^{6}$ | $(8.12^{+0.45}_{-0.43})$ x $10^{-3}$ |
| **gapped G\|A** | (0.64 ± 0.03) x $10^{7}$ | $(2.05^{+0.08}_{-0.08})$ x $10^{-1}$ |
| **gapped G\|C** | (2.05 ± 0.48) x $10^{5}$ | $(2.70^{+0.89}_{-0.67})$ x $10^{-4}$ |
| **gapped G\|G** | (1.14 ± 0.18) x $10^{7}$ | $(6.47^{+1.87}_{-1.45})$ x $10^{-3}$ |
| **gapped G\|U** | (0.45 ± 0.05) x $10^{7}$ | $(4.29^{+0.21}_{-0.20})$ x $10^{-1}$ |
| **gapped U\|A** | (0.47 ± 0.08) x $10^{7}$ | $(4.26^{+0.76}_{-0.64})$ x $10^{-2}$ |
| **gapped U\|C** | (0.39 ± 0.04) x $10^{7}$ | $(2.16^{+0.60}_{-0.47})$ x $10^{-4}$ |
| **gapped U\|G** | (0.57 ± 0.92) x $10^{3}$ | $(9.14^{+28.68}_{-6.93})$ x $10^{-8}$ |
| **gapped U\|U** | (0.56 ± 0.05) x $10^{7}$ | $(5.60^{+0.33}_{-0.31})$ x $10^{-2}$ |

| | | |
|---|---|---|
| **nicked A\|A** | $(0.37 \pm 0.09) \times 10^7$ | $(1.28^{+1.16}_{-0.61}) \times 10^{-4}$ |
| **nicked A\|C** | $(0.50 \pm 0.10) \times 10^7$ | $(1.27^{+0.73}_{-0.46}) \times 10^{-5}$ |
| **nicked A\|G** | $(0.90 \pm 0.15) \times 10^7$ | $(4.23^{+5.14}_{-2.32}) \times 10^{-6}$ |
| **nicked A\|U** | $(1.82 \pm 0.31) \times 10^6$ | $(2.96^{+2.14}_{-1.24}) \times 10^{-5}$ |
| **nicked C\|A** | $(0.57 \pm 0.12) \times 10^7$ | $(6.23^{+3.94}_{-2.42}) \times 10^{-4}$ |
| **nicked C\|C** | $(0.67 \pm 0.10) \times 10^7$ | $(7.75^{+4.46}_{-2.75}) \times 10^{-7}$ |
| **nicked C\|G** | $(0.47 \pm 0.05) \times 10^7$ | $(1.36^{+0.30}_{-0.24}) \times 10^{-5}$ |
| **nicked C\|U** | $(2.96 \pm 0.80) \times 10^6$ | $(8.31^{+0.89}_{-0.81}) \times 10^{-5}$ |
| **nicked G\|A** | $(0.89 \pm 0.19) \times 10^7$ | $(1.36^{+0.69}_{-0.46}) \times 10^{-4}$ |
| **nicked G\|C** | $(0.68 \pm 0.01) \times 10^7$ | $(2.46^{+2.97}_{-1.34}) \times 10^{-6}$ |
| **nicked G\|G** | $(1.14 \pm 0.19) \times 10^7$ | $(1.47^{+1.60}_{-0.76}) \times 10^{-6}$ |
| **nicked G\|U** | $(0.33 \pm 0.05) \times 10^7$ | $(4.19^{+0.53}_{-0.47}) \times 10^{-5}$ |
| **nicked U\|A** | $(0.67 \pm 0.17) \times 10^7$ | $(9.83^{+4.88}_{-3.26}) \times 10^{-4}$ |
| **nicked U\|C** | $(1.00 \pm 0.10) \times 10^7$ | $(1.71^{+0.63}_{-0.46}) \times 10^{-5}$ |
| **nicked U\|G** | $(0.95 \pm 0.15) \times 10^7$ | $(3.61^{+3.23}_{-1.71}) \times 10^{-6}$ |
| **nicked U\|U** | $(0.66 \pm 0.10) \times 10^7$ | $(6.82^{+1.28}_{-1.08}) \times 10^{-4}$ |
| **dangling C\|A, polyA$_{15}$ tail** | $(1.13 \pm 0.08) \times 10^7$ | NA |
| **dangling C\|A** | $(1.93 \pm 0.15) \times 10^7$ | NA |

### 6. Dinucleotide binding

Given the importance of short oligonucleotides in the context of the origin of life, and particularly of imidazolium bridged dinucleotides[4], we measured the binding energy of two canonical dinucleotides to templates containing 2-aminopurine (Supporting Figure 4A) at room temperature.

To perform these experiments, RNA dinucleotides of sequence 5′-GU-3′ and 5′-UG-3′ were hybridized to a dinucleotide gap and the fluorescent emission of 2-aminopurine was monitored over time. The quenching was fitted with a simple hyperbolic binding curve having amplitude and binding constant as free parameters, yielding K equal to 411 μM and 104 μM for 5′-GU-3′ and 5′-UG-3′ respectively (Supporting Figure 4B).

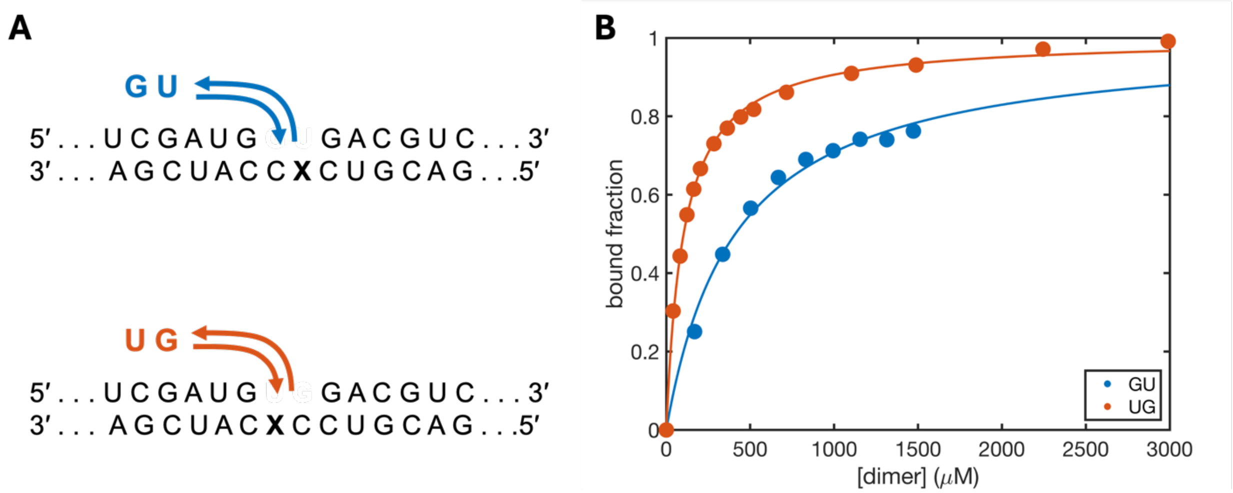

**Supporting Figure S4.** Dinucleotide binding. (**A**) Sketch for experimental design used to measure dinucleotides binding to a dinucleotide gap. The X in the template sequence stands for 2-aminopurine. (**B**) Normalized binding curves as determined through quenching experiments for GU and UG dimers.

The values obtained with predictions using our experimentally determined values for coaxial stacking (see main text) yield $\Delta G_{20°C}$ for 5′-GU-3′ equal to -6.1 kcal/mol and 5′-UG-3′ equal to -7.9 kcal/mol, to be compared with values obtained here equal to -4.5 kcal/mol and -5.3 kcal/mol respectively. The offsets are equal to -1.6 kcal/mol and -2.6 kcal/mol, a difference that can be rationalized as coming from the length dependence discussed in the main text and potentially from an imperfect additivity of the two coaxial interfaces next to the extremely short dinucleotides.

Comparing the binding constants determined this way with $K_M$ values from primer extension experiments from literature in a buffer with comparable ionic strength[5], we found our dissociation constants for 5′-GU-3′ and 5′-UG-3′ to be more stable, with $\Delta\Delta G$ at 20°C equal to 1.9 kcal/mol and 1.2 kcal/mol respectively. We attribute these differences to the presence of the chemically activating imidazolium bridge in the dinucleotides used by Ding et al.[4]

### 7. Predicting binding strength of chemically activated species

Binding strengths of ultrashort (1 ~ 4 nt) chemically activated oligonucleotides are analyzed in this work. The complete dataset of experimentally determined values from literature[6,5,7] and predictions is presented in Supporting Table 6.

**Supporting Table 6. Predictions for binding strength of chemically activated species**

| Species | Experimental System | $\Delta G_{20°C}$ (kcal/mol) Predictions, new coaxial stacking | $\Delta G_{20°C}$ (kcal/mol) Predictions, coaxial stacking from NNDBΩ | $\Delta G_{20°C}$ (kcal/mol) Experiment |
|---|---|---|---|---|
| A*CGC | Upstream Nick | -8.34 | -7.48 | -5.96 |
| A*CG | Upstream Nick | -4.29 | -3.43 | -3.65 |
| A*C | Upstream Nick | -1.48 | -0.62 | -1.74 |
| *A | Upstream Nick | -1.57 | -0.71 | -1.12 |
| *CUGA | Upstream Nick | -7.90 | -7.13 | -7.07 |
| *CGC | Upstream Nick | -7.38 | -6.61 | -7.03 |
| A*A | Double Nick | -2.17 | +1.37 | -4.28 |
| A*C | Double Nick | -3.30 | -0.52 | -3.25 |
| A*G | Double Nick | -5.29 | -2.22 | -5.55 |
| A*U | Double Nick | -1.19 | +1.55 | -2.25 |
| C*A | Double Nick | -4.69 | -1.72 | -4.83 |
| C*C | Double Nick | -5.58 | -3.37 | -4.56 |
| C*G | Double Nick | -6.76 | -4.27 | -6.07 |
| C*U | Double Nick | -3.38 | -1.22 | -2.49 |
| G*A | Double Nick | -5.32 | -1.82 | -5.13 |
| G*C | Double Nick | -6.11 | -3.37 | -3.83 |
| G*G | Double Nick | -8.06 | -5.04 | -6.91 |
| G*U | Double Nick | -3.88 | -1.18 | -2.63 |
| U*A | Double Nick | -2.97 | +1.11 | -3.42 |
| U*C | Double Nick | -3.84 | -0.52 | -2.37 |
| U*G | Double Nick | -5.69 | -2.08 | -4.15 |
| U*U | Double Nick | -1.27 | +2.01 | -1.70 |

### 8. Disentangling hydrogen bond and stacking contributions to the formation of the double helix

To evaluate the effect of hydrogen bonds and coaxial stacking in the transition from single stranded RNA to double stranded RNA, we employed the approach from Zacharias[8]. Briefly, the free energy change associated with the stacking of a fraction *x* of bases is:

$$\Delta G(x) = \Delta G^{stacking} + RTln\left(\frac{x}{1-x}\right)$$

For convenience, Zacharias computes the integral to define a Free Energy function along the *x* coordinates,

$$G(x) = x\Delta G^{stacking} + RT(1-x)ln(1-x) + RTxln(x) + C$$

and uses this to calculate the Free Energy function associated with helix propagation by subtracting the constant term *C* and the NN energy, so that *G(x)* has its minimum along *x* corresponding to the fraction of stacked bases at equilibrium, and it is offset by the energy from the NNDB. For example, for the formation of a CG pair,

$$G(x) = x\Delta G_{CG}^{stacking} + 2RT(1-x)ln(1-x) + 2RTxln(x) - G_{min} - \Delta G_{CG}^{NN,helix\ propagation}$$

where $\Delta G^{stacking}$ is the energy associated with coaxial stacking as determined in this work. This model produces the free energy functions shown in Supporting Figure 5. Given the large energies determined in this work, almost all nucleobases are expected to be stacked at equilibrium (mean $x \approx 0.96$), with helix propagation driven by hydrogen bonds.

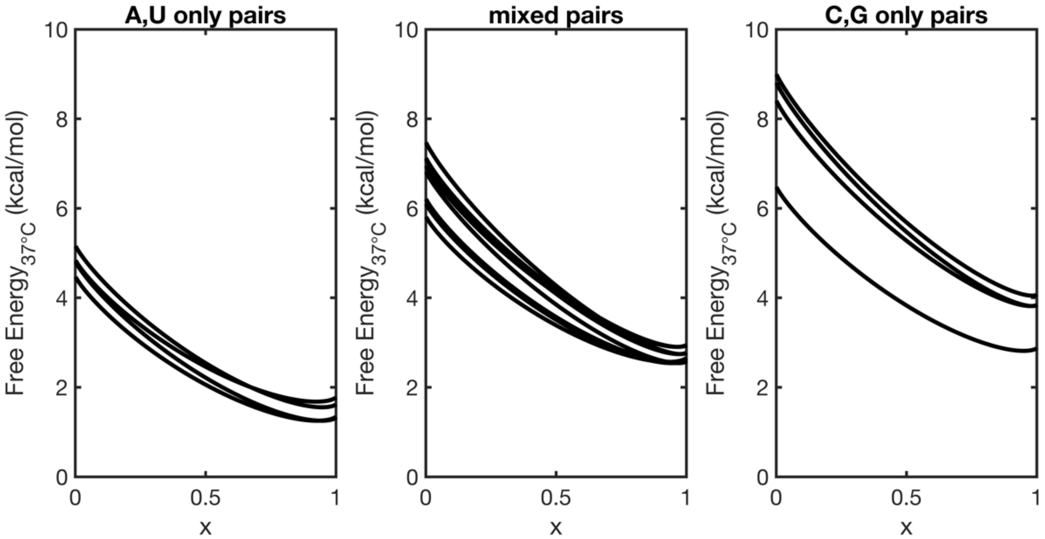

**Supporting Figure S5.** Free energy profile for the formation of a new base pair as a function of the fraction of stacked nucleobases ($x$). The position of the minimum along the $x$ axis corresponds to expected fraction of stacked nucleobases at equilibrium as derived from the experimentally measured coaxial stacking. The Free Energy value at such a minimum corresponds to the value associated with the formation of a given base pair from the NNDB.
